# Age and Social Observation Effects on Theta Synchrony and Its Role in Adolescent Post-Error Control: A Computational Approach

**DOI:** 10.64898/2026.06.29.734887

**Authors:** Felix Zakirov, Kianoosh Hosseini, Ana Garcia Morazzani, Lillian LaPlace, Jeremy W. Pettit, George A. Buzzell

## Abstract

Error monitoring allows for detecting mistakes and adapting behavior. Error monitoring is associated with increased theta (4-7 Hz) EEG activity recorded at midfrontal electrode sites located over the medial frontal cortex. Increases in theta synchrony after errors between midfrontal and lateral electrode sites, located over lateral prefrontal, motor, and sensory/parietal brain regions, are associated with post-error behavioral adaptations. However, there is a lack of research into whether such error-related theta inter-regional synchrony dynamics exhibit age-related changes across adolescence and whether such changes differ as a function of social observation. Moreover, there are discrepancies across studies in terms of how error-related theta synchrony between different electrode sites relates to specific forms of post-error adjustments. In this study, we used behavioral and EEG data from 262 adolescents aged 10-14 years who performed a flanker task twice: alone and while observed by a peer. Regardless of social observation, we found age-related increases in error-related midfrontal-frontolateral and midfrontal-midlateral theta synchrony. Additionally, we found that error-related theta synchrony between midfrontal and posterolateral regions increased only in the alone condition. Leveraging the shrinking spotlight drift diffusion model (SSP-DDM), we identified a positive association between error-related midfrontal-posterolateral theta synchrony and post-error attentional control but found no effects of social observation. These results are the first to demonstrate that error-related theta synchrony increases from early to mid-adolescence. Additionally, this work specifically links error-related midfrontal-posterolateral theta synchrony to post-error attentional control using computational modeling.

## Introduction

Cognitive control refers to a set of mechanisms that allow for monitoring one’s performance and adjusting cognitive/behavioral strategies to achieve task goals (Ullsperger et al., 2014; Yeung, 2014). More specifically, “error monitoring” refers to self-monitoring and detecting when errors occur, as error commission can serve as an alerting signal indicating that “post-error adjustments” to cognition and behavior are needed to avoid future errors. Error monitoring has been associated with enhanced response-locked activity within the theta band recorded at midfrontal electrode sites located over the medial frontal cortex in both youth and adults (Buzzell et al., 2019; Cavanagh & Frank, 2014). Prior EEG work suggests that response-locked theta inter-regional synchrony (i.e., inter-channel phase synchrony; ICPS) between electrodes located over midfrontal and task-relevant cortical regions predicts adaptive changes in behavior on subsequent trials (Buzzell et al., 2019; Cavanagh et al., 2009; Cohen et al., 2009; Cohen & van Gaal, 2013; Driel et al., 2012). However, the evidence for how particular theta dynamics are associated with specific forms of post-error adjustments is inconsistent across studies (Cavanagh et al., 2009; Driel et al., 2012). Moreover, the role of contextual factors (e.g., social observation) in these processes remains unresolved.

Cognitive control develops through childhood and approaches stability across adolescence (Crone & Steinbeis, 2017; Luna, 2009). Adolescents are more sensitive to peer influence than children and adults (Albert et al., 2013; Somerville, 2013; Steinberg & Monahan, 2007), with the effects of social observation on cognitive control potentially changing with age due to the increasing importance of peers across adolescence (Andrews et al., 2021; Somerville, 2013). Buzzell and colleagues (2019) showed that adolescents exhibit similar error-related theta dynamics as adults, including error-related increases in midfrontal theta power and inter-regional synchrony, which were even more pronounced under social observation. However, prior work did not test whether error-related theta dynamics and related social observation effects in adolescents differ with age. Given increases in peer sensitivity and the ongoing maturation of cognitive control systems during adolescence, it is critical to study age effects to understand when and how these changes develop. Thus, the first goal of this study was to investigate the effect of age and social observation (peer observation) on distinct forms of error-related theta inter-regional synchrony across adolescence. A second goal of this study was to investigate the functional role of error-related theta inter-regional synchrony by assessing associations with computational behavioral outcomes, which we describe in further detail below.

Prior work demonstrates that, following the commission of an error, cognitive control systems typically engage post-error adjustments to improve performance in subsequent trials (Danielmeier & Ullsperger, 2011) via at least two mechanisms. One mechanism involves increasing response caution, i.e., a speed-accuracy tradeoff in favor of higher accuracy, which is believed to occur due to motor inhibition (Guan & Wessel, 2022; Ullsperger & Danielmeier, 2016; Wessel, 2018). A second mechanism involves increasing attentional control, i.e., enhancement of task-specific selective attention, which is believed to occur via recruitment of prefrontal and/or parietal regions and their commensurate effects on sensory regions (Cohen & van Gaal, 2013; King et al., 2010; Wessel, 2018). On the behavioral level, these performance adjustments are indexed by a post-error increase in reaction time (RT) (post-error slowing, PES) and accuracy (post-error accuracy, PEA), and reductions in the degree to which RT and/or accuracy are impacted by stimulus congruency (post-error reduction in interference, PERI) (Danielmeier & Ullsperger, 2011). These behavioral measures can be predicted by response-locked theta activity on error trials (Beatty et al., 2020; Buzzell et al., 2019; Cavanagh et al., 2009; Valadez & Simons, 2018). For example, Buzzell and colleagues (2019) have shown that an error-related increase in theta synchrony between midfrontal and lateral-frontal electrodes predicts PERI, suggesting it may reflect recruitment and/or instantiation of attentional control after an error. There is also work suggesting that attentional control may be recruited and/or instantiated via long-range theta synchrony between midfrontal and posterolateral electrodes located over parieto-occipital regions, which has been shown to increase PEA in visual tasks (Cohen et al., 2009; Cohen & van Gaal, 2013). In contrast, error-related synchrony between midfrontal and midlateral electrodes (located over motor regions) has been shown to predict PES (Buzzell et al., 2019) and this may serve as evidence of top-down regulation over motor regions to engage a more cautious post-error response strategy. Still other work shows that error-related theta synchrony between midfrontal and lateral-frontal electrodes located over the dorsolateral prefrontal cortex (DLPFC) also predicts PES (Cavanagh et al., 2009) and is increased when errors are caused by failures of motor control (Driel et al., 2012). Thus, although prior work broadly supports the link between error-related theta inter-regional synchrony and post-error adjustments (i.e., changes in attentional control vs. response caution), discrepancies exist across studies. Moreover, prior work investigating relations between theta inter-regional synchrony and post-error adjustments did not assess the impact of age and social observation across the adolescent period.

One possible reason for the discrepancy across studies is that prior work relied on raw behavioral measures to index post-error adjustments. While raw behavioral measures are useful to assess the outcome of a decision, they lack specificity and make it difficult to assess which stage of cognitive processing (e.g., sensory processing, attention, motor preparation) contributes most to a given behavioral outcome (Ratcliff & McKoon, 2008; White & Kitchen, 2022). In contrast to raw behavioral measures, sequential sampling models—such as drift-diffusion models (DDM)—allow for decomposing the decision-making process into latent components related to different cognitive processes (Boehm et al., 2018; White & Kitchen, 2022). The shrinking spotlight DDM (SSP-DDM, White et al., 2011), reflects a specific DDM that has shown the best fit to the flanker task (White et al., 2011, 2018), commonly used to study error monitoring. Like a standard DDM, the SSP-DDM parameters include non-decision time (*Ter*), comprising the time associated with stimulus encoding and response execution (Dutilh et al., 2019; Voss et al., 2004), and boundary separation (*a*), which reflects the amount of evidence needed to initiate a given response, thus indicating response caution. Crucially, to account for a gradual narrowing of attention in the flanker task, SSP-DDM introduces three additional parameters: perceptual strength (*p*), the width of the initial attentional window (*sd_a_*), and the speed of narrowing of the attentional window (shrinking rate, *r_d_*). After model fitting, it is recommended that the latter two parameters be converted into a single *sd_a_*/*r_d_* ratio metric, which represents attentional control (White et al., 2018). Therefore, we employed the SSP-DDM to carefully distinguish between two distinct forms of post-error adjustments: attentional control (*sd_a_*/*r_d_*ratio) and response caution (*a* boundary separation). Thus, the second goal of this study was to investigate how distinct forms of error-related theta inter-regional synchrony, as a function of age and social observation, relate to specific forms of post-error adjustments.

In this study, we first characterized age-related changes in adolescent error-related theta inter-regional synchrony and tested the impact of social observation. Given the ongoing development of error monitoring during adolescence and the notion that adolescents are more sensitive to peer influence, we expected that theta synchrony responses would increase with age and be enhanced under social observation. Further, based on the notion that error-related theta inter-regional synchrony instantiates post-error adjustments (attentional control and response caution), we leveraged the SSP-DDM to investigate the functional significance of error-related theta synchrony and its potential age-related and observation-related changes.

## Materials and Methods

### Participants

Participants were 262 adolescents ages 10 to 14 years (M = 12.17 years, SD = 1.04; 142 f, 120 m; 195 White, 27 Black, 6 Asian, 2 American Indian, 14 more than one race, 18 unknown race; 189 Hispanic/Latino, 70 other, 3 unknown ethnicity) who completed the in-person baseline assessment of a larger, ongoing longitudinal study. Only data from the baseline assessment wave are used in the current report. To be included, participants were required not to have an intellectual disability, psychotic disorder, or substance use disorder and not be taking short-acting psychopharmacological medications (e.g., benzodiazepines). Prior to the study, written informed consents were obtained from all parents, and assents were obtained from the children.

Out of the initial 262 participants, three participants were excluded due to experimental inconsistencies/error. Data from four participants (two dyads) in the social (observed) condition were also excluded because they later reported being acquainted with their dyad partners prior to the study. As a result of discomfort while wearing the EEG cap, EEG data were not obtained from two participants in both conditions and from one participant only in the non-social condition. Additionally, EEG data of one participant was excluded from both conditions due to an underlying condition that may have compromised EEG data quality. For each condition, EEG and behavioral data were excluded from further analyses for any participants in which flanker task accuracy was < 60% or for which missed responses were > M + 3 SD. Further data removal due to insufficient behavior or EEG trial counts (see below) yielded 224 participants with valid behavioral and/or EEG data for the non-social condition and 227 participants for the social condition. Further details regarding condition-wise (social/non-social) and trial type-wise (error/correct) data availability following data cleaning are provided for each data type below and in Tables 1-2.

**Table 1.**
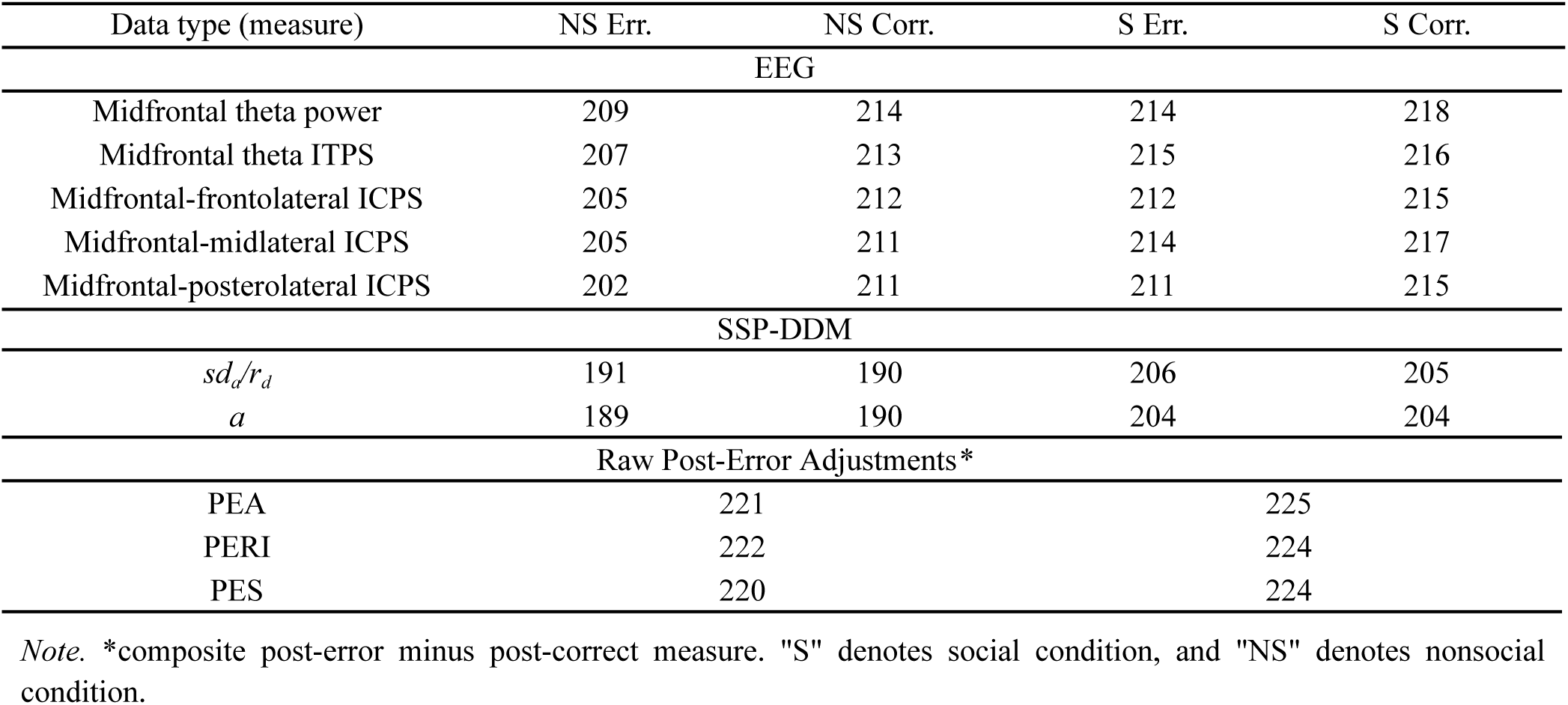
Number of Participants Included in Each Analysis per Data Type per Measure.

**Table 2.** Number of Trials/Epochs (M ± SD) Included in Each Analysis per Data Type per Participant.

| Data type (measure) | NS Err. | NS Corr. | S Err. | S Corr. |
| --- | --- | --- | --- | --- |
| EEG | 34.39 ± 15.74 | 139.31 ± 25.8 | 34.61 ± 15.15 | 139.18 ± 25.54 |
| SSP-DDM/Post-Error Behavior | 37.91 ± 13.23 | 138 ± 17.27 | 37.3 ± 13.57 | 139.73 ± 16.2 |

*Note.* "S" denotes social condition, and "NS" denotes nonsocial condition.

### Study Visit

Two participants typically visited the laboratory at the same time, forming dyads. In instances when only one participant could attend in person, a remote community volunteer was assigned to serve as a dyad partner (15.4% of sessions). Participants within a dyad were age-matched within ± 1.5 years of each other; the biological sex of the dyad partners varied randomly. During the visit, each participant in the dyad completed a demographics survey as part of a larger battery of surveys. Subsequently, EEG was recorded, and participants performed the arrow flanker task (Figure 1) in two conditions: social and non-social in a counterbalanced order. In the social condition, the first participant was informed that the other participant (dyad partner) would observe them and read the feedback aloud via a video call on an iPad (Apple Inc.). For more details about the social interaction protocol, see Supplementary Materials.

**Figure 1.**
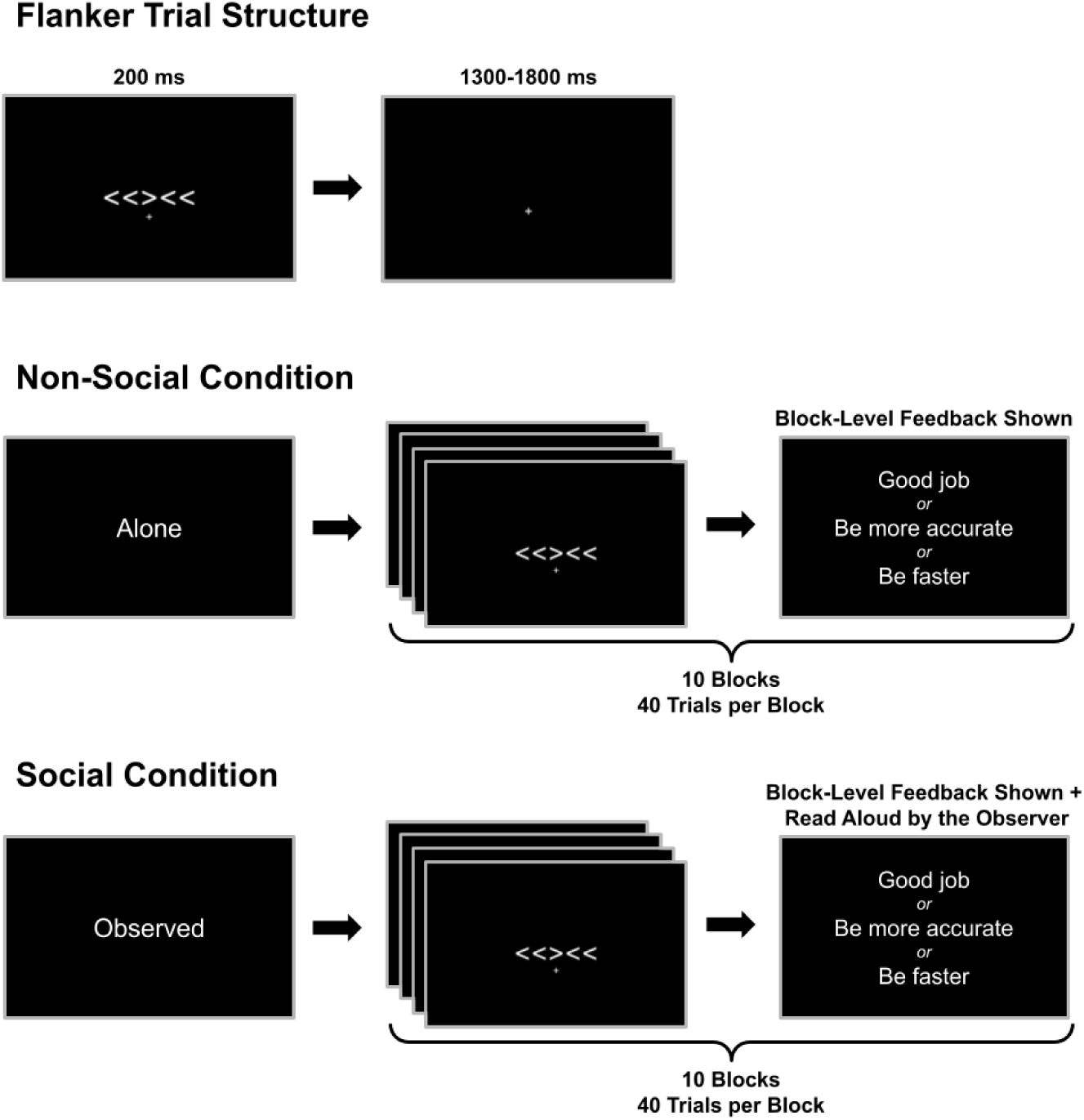
Flanker Task Structure. *Note.* All 10 blocks of each condition (400 trials total) were completed consecutively, with no alternation between the two conditions. The order of conditions was counterbalanced across participants.

### Task

Participants completed a modified version (Buzzell et al., 2017, 2019) of the flanker task (Eriksen & Eriksen, 1974) twice: once in a non-social condition and once in a social condition (Figure 1), in counterbalanced order. In the non-social condition, participants performed the task alone; in the social condition, they performed the task while being observed by a peer via Zoom (Zoom Communications Inc.) video call. Each condition consisted of 10 blocks, 40 trials each, and participants were able to practice before the main task. In each trial of the task, a central arrowhead, pointing to the left or right, was presented flanked by two additional arrowheads on each side pointing in the same (congruent trials) or the opposite (incongruent trials) direction as the central arrowhead. Each arrowhead had a height and width of 0.6 degrees of visual angle. In each trial, the arrows were presented for 200 ms and then were followed by a response window that varied randomly from 1300 to 1800 ms which also served as an intertrial interval (ITI). A fixation cross remained on the screen throughout all trials within a block (40 trials). All 10 blocks of each condition (400 trials total) were completed consecutively, with no alternation between the two conditions. Congruent and incongruent trials were presented in random order and in equal proportion. Participants were instructed to indicate the direction of the middle arrow by pressing the left/right button of a button box (The Black Box ToolKit Ltd., Sheffield, UK) using their left/right thumb, respectively. Participants were instructed to respond as quickly and as accurately as they could. To maintain sufficient error rates, feedback was presented onscreen after each block (and additionally read by the peer in the social condition): “Be more accurate” for < 75% accuracy, “Respond faster” for > 90% accuracy, and “Good job” for accuracy within 75%-90%. A reminder of task instructions was provided on the computer screen before each block. Stimuli were presented on an ASUS VG248QG 24” 165 Hz screen, and participants were seated 80 cm from the screen. The lights were dimmed in the study room, and the experimenter was not present in the study room while participants completed the task.

### EEG Acquisition and Preprocessing

EEG was recorded using a 64-channel Brain Vision actiCHamp Plus amplifier, coupled with an actiCap electrode cap with a custom layout (see Figure 2), and managed through Recorder software (Brain Products GmbH, Gilching, Germany). The data was sampled at a rate of 1000 Hz and recorded employing a vertex reference and frontocentral ground configuration. To ensure optimal recording conditions, impedances were adjusted to levels below 25 KΩ before initiating the recording session. See Tables 1-2 for the resulting numbers of participants and epochs included in the analyses for each measure.

**Figure 2.**
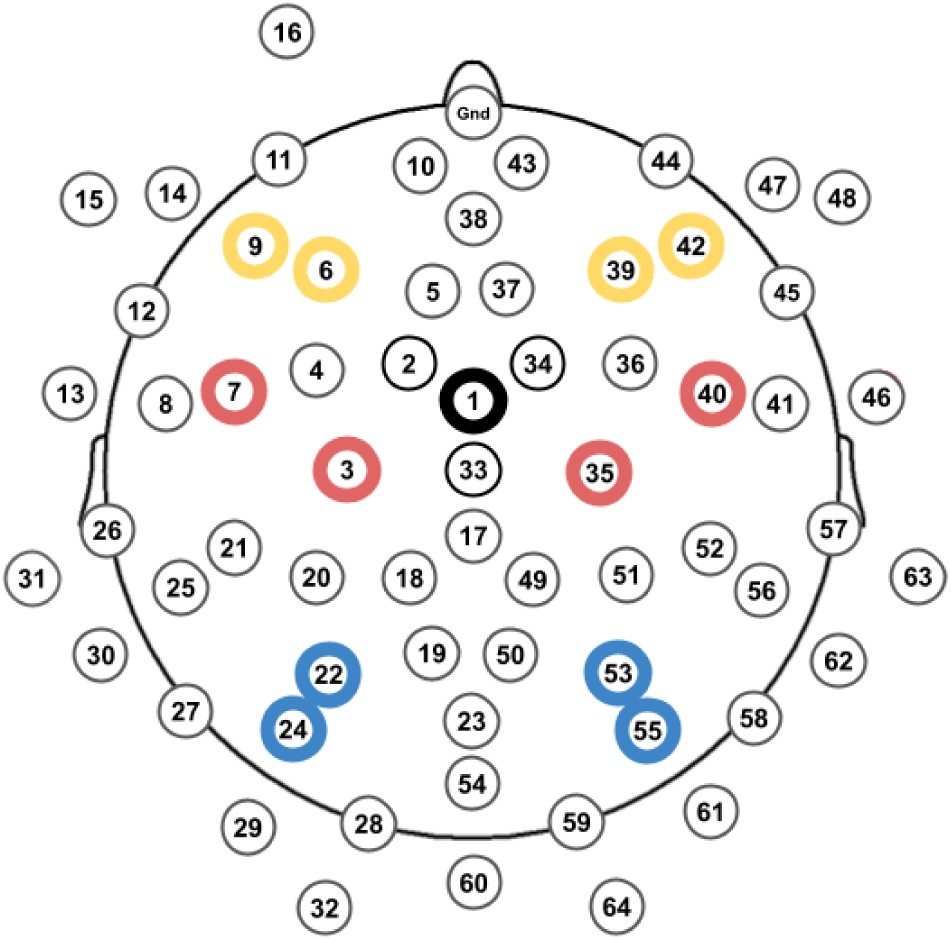
Custom 64-Channel Layout and Electrode Clusters Included in the Analysis. *Note.* Black: midfrontal seed electrode for computing ICPS measures; yellow: frontolateral cluster (located over lateral prefrontal sites); red: midlateral cluster (located over central motor and pre-motor sites); blue: posterolateral cluster (located over occipitoparietal sites).

EEG processing and analysis were performed in MATLAB (MathWorks Inc., Sherborn, MA, USA) using the EEGLAB toolbox (Delorme & Makeig, 2004) and custom MATLAB scripts. Preprocessing was performed following a modified version of the Maryland Analysis of Developmental EEG (MADE) pipeline (Debnath et al., 2020); see Supplementary Materials for further details. For time-frequency (TF) based measures, the data were epoched from -1000 to 2000 ms relative to the response. Prior to computing the TF measures, a Laplacian transform was applied to the EEG data to improve the spatial specificity of subsequent analyses. A Laplacian transform acts as a spatial filter that attenuates low spatial frequencies and highlights high spatial frequencies (local activity at each electrode site), thus minimizing the effects of volume conduction and improving the interpretation of inter-regional synchrony measures (Carvalhaes & de Barros, 2015; Kayser & Tenke, 2015).

Prior to TF decomposition, individual trials were removed if RT was < 150 ms (premature responses) or if multiple responses were given. All TF measures for each trial type (error/correct) in each condition (social/non-social) were computed only for participants with at least six valid epochs, as this has been shown to be necessary for a reliable estimate of response-locked activity (Pontifex et al., 2010; Steele et al., 2016). Unless specified otherwise, data from only incongruent trials were used for computing TF measures and statistical analyses (error versus correct) in order to isolate error-related effects and remove the confound of accuracy with congruency (Buzzell et al., 2019).

TF decomposition was performed via Morlet wavelet convolution for frequencies between 1-30 Hz divided into 59 linearly spaced steps. To balance frequency and temporal precision, the number of wavelet cycles was linearly increased as the frequency increased, starting from 3 cycles at 1 Hz to 10 cycles at 30 Hz (Morales & Bowers, 2022). All TF measures were computed separately for each type of epoch for accuracy (error/correct) and condition (social/non-social). To assess response-locked theta inter-regional synchrony, our primary measure of interest, inter-channel phase synchrony (ICPS), was computed as the average of phase angle differences across trials between electrodes and could vary from 0 to 1. ICPS was computed between a seed midfrontal electrode and three separate lateral regions: 1) bilateral frontolateral electrode clusters; 2) bilateral midlateral electrode clusters; 3) bilateral posterolateral electrode clusters (see Figure 2). ICPS was computed between individual electrodes and then averaged within clusters. To account for the unequal number of trials in ICPS analysis, a subsampling algorithm was used, such that for each condition, six random trials were sampled 100 times, and then results were averaged. Following TF decomposition, data were decibel-corrected relative to a -400 to -200 ms pre-response baseline. In addition to ICPS, we also computed response-locked theta power and intertrial phase synchrony (ITPS) over the midfrontal electrode cluster for preliminary/supplementary analyses (see Supplementary Materials).

For subsequent statistical analyses, ICPS for each condition (social/non-social) and trial type (error/correct), were extracted and then averaged within a 4-7 Hz and 0-250 ms region of interest (ROI). Additionally, the ICPS scores were collapsed across hemispheres for parsimony in the main analysis.

### DDM

The SSP-DDM (White et al., 2011) was used to model behavioral RT and accuracy data. In contrast to a traditional DDM, the SSP-DDM contains additional parameters accounting for the dynamics of top-down attention in visual-spatial conflict tasks, such as the flanker task. Specifically, the drift rate (*v)* parameter of the SSP-DDM considers the perceptual strength (*p)* of target and distractor stimuli, the initial size of the attentional window (*sd_a_*), and the shrinking rate (*r_d_*) of that window. As a measure of attentional control, *sd_a_*/*r_d_*ratio was computed from the resulting parameter values obtained after the fitting. For subsequent analyses, the *sd_a_*/*r_d_* ratio was reversed so that higher values corresponded to an increase in attentional control and vice versa for lower values. Similar to a classic DDM, the SSP-DDM also includes the following additional parameters: boundary separation *(a)*, indicating response caution (higher values corresponding to increased response caution); nondecision time *(Ter),* indicating the duration of stimulus encoding and motor processing; and starting point *(z)*, indicating the initial bias towards a specific response. In the current study, the starting point was fixed at the middle point (*a*/2), assuming no systematic bias towards any choice option (equal proportion of congruent/incongruent and left/right trials). The description of the model fitting procedure is provided in the Supplementary Materials.

Prior to the SSP-DDM fitting, behavioral data for social and non-social conditions were processed separately. Individual trials were removed if RT was < 150 ms (premature responses) or if multiple responses were given. Further, to subset post-error and post-correct trials for modeling, only trials following incongruent trials were used to remove the confound of stimulus congruency (Buzzell et al., 2019). Based on the results of a parameter recovery simulation (see Supplementary Materials and Figure S1) a minimum of 16 post-error trials were determined as being sufficient for SSP-DDM modeling and parameter recovery. Thus, participants with fewer than 16 post-error trials per condition were excluded from model fitting, resulting in the removal of 21 participants from the social condition and 32 from the non-social condition for SSP-DDM modeling.

Following model fitting, participants with unusually high fit statistics (χ^2^ > 200) for a given condition (social/non-social) or trial type (post-error/post-correct) were excluded from further analyses. See Tables 1-2 for the resulting numbers of individual trials and participants included in the analyses for each measure.

### Analytic Strategy

Prior to computing statistics, outliers were removed from all measures (± 3 SD), separately for each condition (social/non-social) and trial type (error/correct); see Table 1 for the number of participants included in the analysis of each measure. For all statistical analyses, we used linear mixed-effects models (LMMs), which capture hierarchical individual variability and are robust to missing data (Heise et al., 2022). As described in further detail within the subsequent section, models were fit to investigate the fixed effects of accuracy, age, and social observation. Biological sex (male/female), as well as whether the dyad partner was in-person/remote, were also included in all models as fixed effects to control for these variables. Participant was included as a random effect (intercept) in all models. Sum contrast coding (−1/1) was applied to all categorical variables, and all continuous variables were z-transformed prior to model fitting. The lme4 package (Bates et al., 2015) in R (R Core Team, 2021) was used to fit LMMs, and effects were estimated via restricted maximum likelihood (REML; Patterson & Thompson, 1971). Follow-up interaction tests were conducted using the emmeans package (Lenth, 2025), and a false discovery rate (FDR) correction (Benjamini & Hochberg, 1995) was applied to all *p* values. A detailed description of the statistical models for each of the analyses reported in the main text is provided below. We additionally provide a description of preliminary flanker task behavior (Tables S2-S3) and midfrontal theta activity (power and ITPS; Tables S4-S5) analyses, as well as complementary analyses with raw post-error behavior measures (Tables S11-S22) in the Supplementary Materials.

#### Effects of Accuracy, Age, and Social Observation on Theta ICPS

In the primary analyses, we first sought to test whether and how error-related theta ICPS differs as a function of accuracy, social observation, and age. An LMM was separately fit for each of the three theta ICPS measures (midfrontal-frontolateral, midfrontal-midlateral, and midfrontal-posterolateral; see Figure 2). In each model, accuracy (correct/incorrect), condition (social/non-social), age in months, and their interactions, were included as fixed effects. We report the results of these three statistical models in the main text.

#### Investigating the Functional Role of Error-Related Theta ICPS in Post-Error Control

Having tested the effects of accuracy, age, and social observation on region-specific error-related theta ICPS measures, we next sought to examine the functional significance of these neural measures and their age-related and observation-related changes by examining associations with post-error SSP-DDM measures of attentional control (*sd_a_*/*r_d_*) and response caution (*a*). SSP-DDM parameters (*a*, *sd_a_*/*r_d_*) and ICPS predictors were entered into statistical models as difference scores (post-error minus post-correct, and error minus correct, respectively). We fit two statistical models predicting either post-error attentional control (*sd_a_*/*r_d_*) or response caution (*a),* with error-related theta ICPS measures (midfrontal-frontolateral, midfrontal-midlateral, and midfrontal-posterolateral), and their interactions with age and social observation, entered as fixed effects. Based on prior literature (Buzzell et al., 2019; Cavanagh et al., 2009; Cohen et al., 2009; Cohen & van Gaal, 2013; Driel et al., 2012), we hypothesized that midfrontal-frontolateral and/or midfrontal-posterolateral theta ICPS would predict post-error attentional control (*sd_a_/r_d_*), whereas midfrontal-midlateral theta ICPS would predict response caution (*a*). Nonetheless, inclusion of all three ICPS measures within the same statistical models retained parsimony while also allowing us to control for the effects of each ICPS measure simultaneously. We report the full results and outputs of these two statistical models in the main text. For completeness, we also fit a more exhaustive set of exploratory models in which we predicted each of the two SSP-DDM measures (*sd_a_*/*r_d_*and *a*) using each of the three theta ICPS measures entered as predictors within separate models (six models total). All other fixed effects and interactions in these exploratory models were otherwise identical to the primary models described above. These six exploratory models revealed no qualitative differences compared to the full models, and we report the full outputs of these exploratory models in the Supplementary Materials (Tables S5-S10).

## Results

### Effects of Accuracy, Age, and Social Observation on Theta ICPS

We investigated the effects of accuracy, age, and social observation, as well as their interaction on error-related theta ICPS (inter-regional synchrony) using separate LMMs (Table 3) for each electrode cluster of interest (Figure 2). A detailed depiction of the results for each ICPS measure, as well as topographies and surface plots (Figure 3), is provided below.

**Table 3.** Results of Statistical Models for Effects of Accuracy, Age, and Social Observation on Theta ICPS.

| Parameter | Midfrontal-Frontolateral |  |  |  |  |  | Midfrontal-Midlateral |  |  |  |  |  | Midfrontal-Posterolateral |  |  |  |  |  |
| --- | --- | --- | --- | --- | --- | --- | --- | --- | --- | --- | --- | --- | --- | --- | --- | --- | --- | --- |
| | $\beta$ | SE | df | t | p | 95% CI | $\beta$ | SE | df | t | p | 95% CI | $\beta$ | SE | df | t | p | 95% CI |
| (Intercept) | -0.02 | 0.06 | 227.50 | -0.39 | .697 | [-0.13, 0.09] | -0.02 | 0.05 | 216.87 | -0.46 | .646 | [-0.12, 0.08] | 0.02 | 0.06 | 218.31 | 0.32 | .749 | [-0.10, 0.14] |
| Accuracy | -0.45 | 0.03 | 614.84 | -17.25 | < .001*** | [-0.51, -0.40] | -0.58 | 0.02 | 611.41 | -24.61 | < .001*** | [-0.63, -0.53] | -0.40 | 0.03 | 609.37 | -14.54 | < .001*** | [-0.45, -0.34] |
| Condition | -0.02 | 0.03 | 650.94 | -0.63 | .528 | [-0.07, 0.04] | 0.01 | 0.02 | 645.78 | 0.55 | .583 | [-0.03, 0.06] | 0.00 | 0.03 | 641.57 | 0.17 | .864 | [-0.05, 0.06] |
| Age | 0.20 | 0.04 | 227.20 | 5.18 | < .001*** | [0.12, 0.27] | 0.18 | 0.04 | 218.54 | 5.20 | < .001*** | [0.11, 0.25] | 0.07 | 0.04 | 221.85 | 1.59 | .112 | [-0.01, 0.15] |
| Sex | 0.02 | 0.04 | 222.54 | 0.56 | .575 | [-0.05, 0.10] | -0.02 | 0.04 | 215.92 | -0.43 | .664 | [-0.08, 0.05] | -0.07 | 0.04 | 219.24 | -1.66 | .098 | [-0.15, 0.01] |
| Peer mode | 0.04 | 0.06 | 226.55 | 0.64 | .520 | [-0.07, 0.14] | 0.05 | 0.05 | 216.02 | 0.95 | .341 | [-0.05, 0.15] | -0.01 | 0.06 | 217.45 | -0.21 | .837 | [-0.13, 0.10] |
| Accuracy * Condition | 0.03 | 0.03 | 615.05 | 1.28 | .202 | [-0.02, 0.09] | -0.02 | 0.02 | 608.76 | -0.75 | .453 | [-0.06, 0.03] | 0.00 | 0.03 | 605.67 | 0.11 | .912 | [-0.05, 0.06] |
| Accuracy * Age | -0.09 | 0.03 | 615.19 | -3.30 | .001** | [-0.14, -0.04] | -0.11 | 0.02 | 612.35 | -4.66 | < .001*** | [-0.16, -0.06] | 0.00 | 0.03 | 607.43 | 0.00 | .997 | [-0.05, 0.05] |
| Condition * Age | -0.02 | 0.03 | 659.68 | -0.63 | .530 | [-0.07, 0.04] | 0.02 | 0.02 | 651.26 | 0.67 | .502 | [-0.03, 0.06] | -0.04 | 0.03 | 649.76 | -1.57 | .116 | [-0.10, 0.01] |
| Accuracy * Condition * Age | 0.01 | 0.03 | 618.30 | 0.45 | .654 | [-0.04, 0.06] | 0.01 | 0.02 | 609.14 | 0.22 | .823 | [-0.04, 0.05] | 0.06 | 0.03 | 605.95 | 2.09 | .037* | [0.00, 0.11] |

*Note.* Significance codes: \* p < .05, \*\* p < .01, \*\*\* p < .001. Degrees of freedom for the fixed effects were estimated using Satterthwaite's approximation.

**Figure 3.**
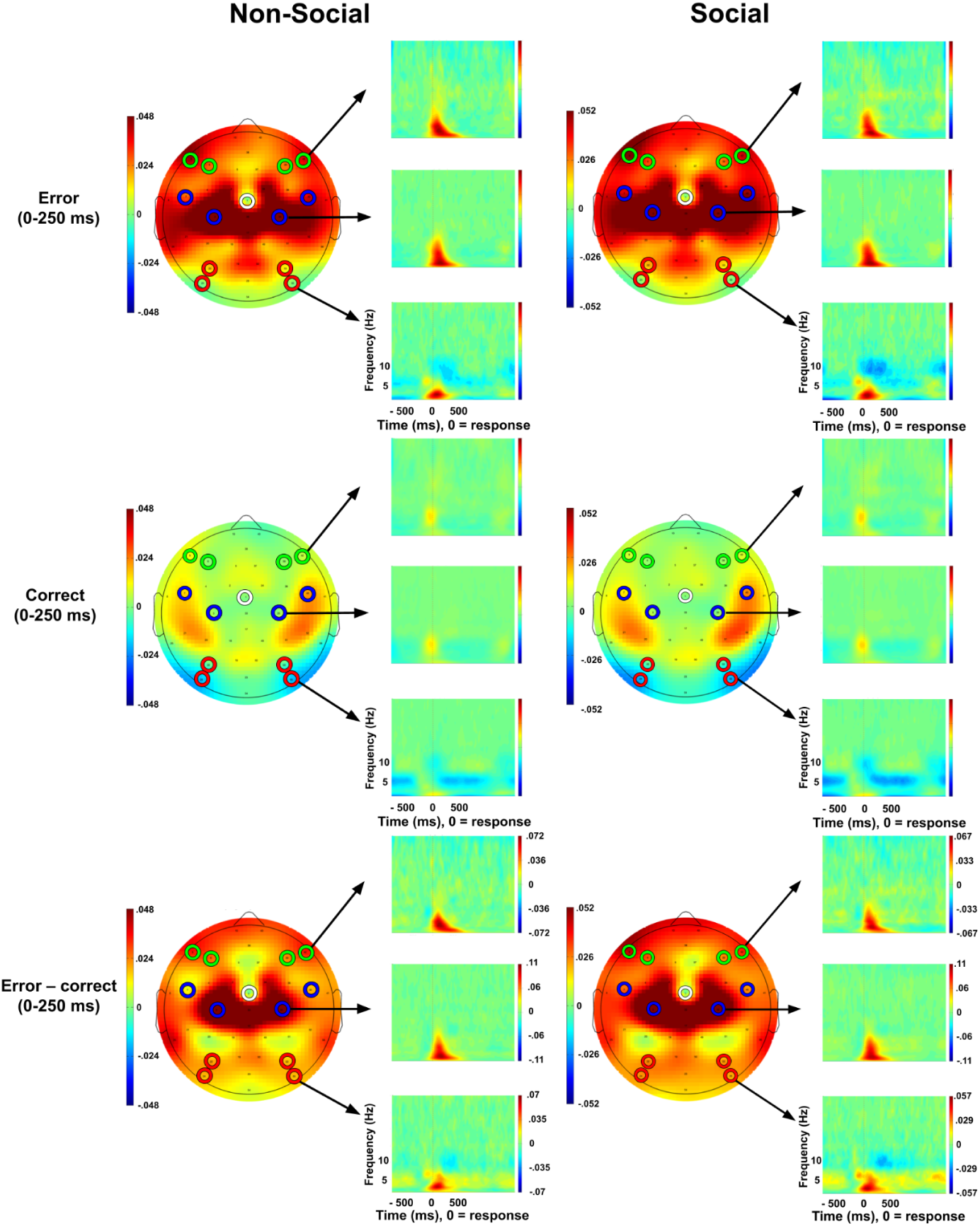
Theta Inter-Regional Synchrony Between Midfrontal and Lateral Scalp Regions. *Note.* Response-locked theta ICPS seeded at a midfrontal electrode (shown in white) in the social and non-social conditions, separately for error, correct, and the error-correct difference. Bilateral frontolateral, midlateral, and posterolateral clusters are shown in yellow, red, and blue, respectively.

### Midfrontal-Frontolateral ICPS

The LMM predicting midfrontal-frontolateral ICPS (Table 3) revealed a significant main effect of accuracy, with greater ICPS values for error compared to correct responses (β = -0.45, 95% CI [-0.51, -0.40], p < .001), as well as a significant main effect of age, such that increased age was associated with greater ICPS (β = 0.20, 95% CI [0.12, 0.27], p < .001). Furthermore, there was a significant accuracy * age interaction (β = -0.09, 95% CI [-0.14, -0.04], p = .001) (Figure 4A). Follow-up simple slopes analyses showed that the positive effect of age was significant for both error-related ICPS (β = 0.29, 95% CI [0.19, 0.38], p < .001) and correct-related ICPS (β = 0.11, 95% CI [0.02, 0.20], p = .017), although it was significantly stronger for error-related ICPS (β = 0.18, 95% CI [0.07, 0.28], p = .001). There were no significant effects involving condition (all p >= .202; Table 3).

**Figure 4.**
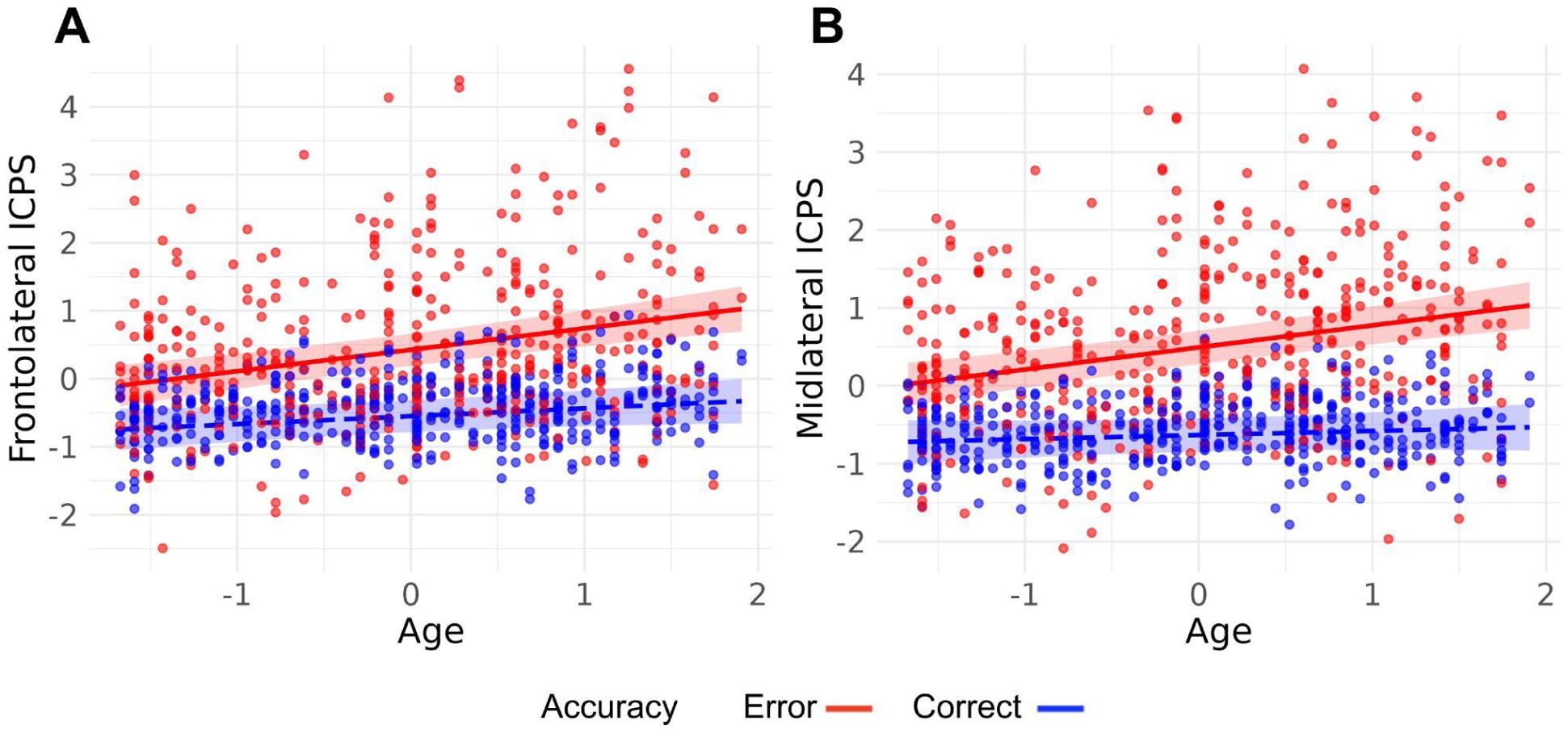
Effects of Age and Response Accuracy on Theta Inter-Regional Synchrony. *Note.* Age-related increases in (A) midfrontal-frontolateral and (B) midfrontal-midlateral theta ICPS. Age-related increases in error-related responses are significantly greater than in correct-related responses. Shaded regions represent 95% confidence intervals.

#### Midfrontal-Midlateral ICPS

The LMM predicting midfrontal-midlateral ICPS (Table 3) exhibited similar patterns, with a significant main effect of accuracy, with increased ICPS values for error compared to correct responses (β = -0.58, 95% CI [-0.63, -0.53], p < .001), and a significant main effect of age, such that increased age was associated with greater ICPS (β = 0.18, 95% CI [0.11, 0.25], p < .001). Additionally, there was a significant accuracy * age interaction (β = -0.11, 95% CI [-0.16, -0.06], p < .001) (Figure 4B). Follow-up simple slopes analyses showed that the positive effect of age was significant only for error-related ICPS (β = 0.29, 95% CI [0.21, 0.38], p < .001) but not for correct-related ICPS (β = 0.07, 95% CI [-0.01, 0.16], p = .084). There were no significant effects involving condition (all p >= .453; Table 3).

#### Midfrontal-Posterolateral ICPS

The LMM predicting midfrontal-posterolateral ICPS (Table 3) revealed a significant main effect of accuracy, with greater ICPS values for error compared to correct responses (β = -0.40, 95% CI [-0.45, -0.34], p < .001). Additionally, there was a significant accuracy * age * condition interaction (β = 0.06, 95% CI [0.00, 0.11], p = .037) (Figure 5). Follow-up analyses probing this interaction revealed a significant age-related increase in error-related ICPS only in the non-social condition (β = 0.17, 95% CI [0.02, 0.31], p = .018), with no corresponding age effect for error-related ICPS in the social condition (β = -0.04, 95% CI [-0.18, 0.11], p = .578), nor for correct-related ICPS in either the non-social condition (β = 0.05, 95% CI [-0.09, 0.19], p = .402) or the social condition (β = 0.08, 95% CI [-0.06, 0.22], p = .411). Critically, we also found that the simple slopes for the effect of age on error-related ICPS in the social vs. nonsocial conditions significantly differed from each other (β = -0.20, 95% CI [-0.36, -0.05], p = .010).

**Figure 5.**
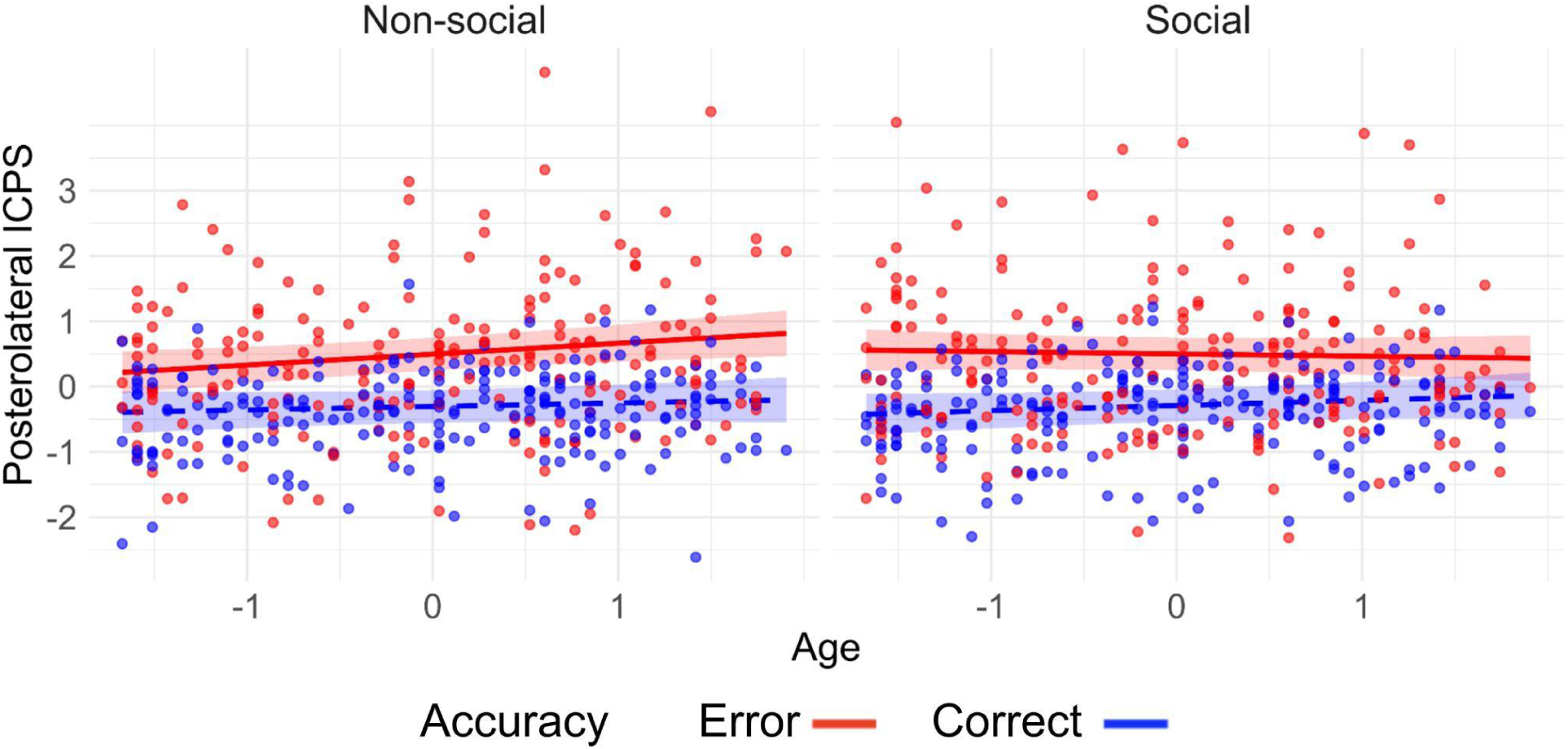
Age-Related Increase in Midfrontal-Posterolateral Theta ICPS Conditional on Response Accuracy and Social Observation. *Note.* Age-related increases in error-related responses were significant only in the non-social condition. Shaded regions represent 95% confidence intervals.

### Role of Error-Related Theta ICPS in Post-Error Adjustments

Having tested the effects of accuracy, age, and social observation on error-related theta ICPS, we then sought to clarify the functional role of these neural measures by examining associations with measures of post-error control. We tested links between error-related theta ICPS and two forms of post-error control—attentional control and response caution—using measures derived from the SSP-DDM (*sd_a_/r_d_*and *a*). Below, we present detailed results and full model outputs (Tables 4-5).

**Table 4.** Error-Related Theta ICPS Predicting SSP-DDM Attentional Ratio (sd_a_/r_d_) Model Statistics.

| Parameter | $\beta$ | SE | df | t | p | 95% CI |
| --- | --- | --- | --- | --- | --- | --- |
| (Intercept) | -0.06 | 0.08 | 193.59 | -0.77 | .443 | [-0.23, 0.10] |
| ICPS frontolateral | -0.03 | 0.07 | 325.31 | -0.35 | .724 | [-0.16, 0.11] |
| Condition | -0.05 | 0.05 | 167.94 | -0.98 | .330 | [-0.16, 0.05] |
| Age | -0.01 | 0.06 | 177.22 | -0.17 | .865 | [-0.12, 0.10] |
| ICPS midlateral | -0.01 | 0.07 | 303.78 | -0.18 | .857 | [-0.15, 0.13] |
| ICPS posterolateral | 0.17 | 0.06 | 309.77 | 2.71 | .007** | [0.05, 0.29] |
| Sex | -0.02 | 0.06 | 174.77 | -0.42 | .678 | [-0.13, 0.09] |
| Peer mode | 0.07 | 0.08 | 194.40 | 0.82 | .415 | [-0.09, 0.23] |
| ICPS frontolateral * Condition | -0.10 | 0.07 | 309.93 | -1.42 | .156 | [-0.24, 0.04] |
| ICPS frontolateral * Age | 0.01 | 0.07 | 325.97 | 0.07 | .941 | [-0.14, 0.15] |
| Condition * Age | -0.00 | 0.05 | 169.41 | -0.09 | .928 | [-0.11, 0.10] |
| Condition * ICPS midlateral | 0.12 | 0.07 | 266.43 | 1.73 | .085 | [-0.01, 0.26] |
| Age * ICPS midlateral | 0.08 | 0.07 | 309.45 | 1.15 | .251 | [-0.05, 0.21] |
| Condition * ICPS posterolateral | 0.04 | 0.06 | 283.71 | 0.68 | .498 | [-0.08, 0.16] |
| Age * ICPS posterolateral | -0.01 | 0.06 | 322.05 | -0.15 | .879 | [-0.13, 0.11] |
| ICPS frontolateral * Condition * Age | 0.03 | 0.07 | 314.46 | 0.40 | .690 | [-0.11, 0.17] |
| Condition * Age * ICPS midlateral | -0.01 | 0.07 | 281.50 | -0.12 | .903 | [-0.14, 0.12] |
| Condition * Age * ICPS posterolateral | -0.04 | 0.06 | 302.14 | -0.63 | .528 | [-0.16, 0.08] |

*Note.* Significance codes: \* $p < .05$ , \*\* $p < .01$ , \*\*\* $p < .001$ . Degrees of freedom for the fixed effects were estimated using Satterthwaite's approximation.

**Table 5.** Error-Related Theta ICPS Predicting SSP-DDM Boundary Separation (a) Model Statistics.

| Parameter | $\beta$ | SE | df | t | p | 95% CI |
| --- | --- | --- | --- | --- | --- | --- |
| (Intercept) | 0.02 | 0.08 | 190.92 | 0.30 | .763 | [-0.13, 0.18] |
| ICPS frontolateral | 0.03 | 0.07 | 320.23 | 0.35 | .726 | [-0.11, 0.16] |
| Condition | 0.01 | 0.05 | 167.37 | 0.18 | .855 | [-0.09, 0.11] |
| Age | 0.11 | 0.06 | 170.91 | 1.92 | .056 | [0.00, 0.22] |
| ICPS midlateral | -0.02 | 0.07 | 292.26 | -0.33 | .742 | [-0.16, 0.11] |
| ICPS posterolateral | 0.11 | 0.06 | 302.63 | 1.74 | .082 | [-0.01, 0.22] |
| Sex | 0.13 | 0.06 | 170.40 | 2.32 | .021* | [0.02, 0.24] |
| Peer mode | 0.02 | 0.08 | 192.93 | 0.29 | .771 | [-0.13, 0.18] |
| ICPS frontolateral * Condition | 0.06 | 0.07 | 313.75 | 0.87 | .385 | [-0.07, 0.20] |
| ICPS frontolateral * Age | -0.15 | 0.07 | 321.35 | -2.03 | .044* | [-0.29, -0.01] |
| Condition * Age | 0.06 | 0.05 | 167.23 | 1.06 | .293 | [-0.05, 0.16] |
| Condition * ICPS midlateral | -0.07 | 0.07 | 270.79 | -0.98 | .329 | [-0.20, 0.07] |
| Age * ICPS midlateral | -0.08 | 0.07 | 298.68 | -1.22 | .225 | [-0.21, 0.05] |
| Condition * ICPS posterolateral | 0.01 | 0.06 | 290.62 | 0.13 | .897 | [-0.11, 0.12] |
| Age * ICPS posterolateral | 0.09 | 0.06 | 314.71 | 1.46 | .145 | [-0.03, 0.21] |
| ICPS frontolateral * Condition * Age | -0.09 | 0.07 | 315.78 | -1.17 | .241 | [-0.23, 0.05] |
| Condition * Age * ICPS midlateral | 0.05 | 0.07 | 282.68 | 0.82 | .412 | [-0.07, 0.18] |
| Condition * Age * ICPS posterolateral | -0.03 | 0.06 | 305.08 | -0.43 | .665 | [-0.14, 0.09] |

*Note.* Significance codes: \* $p < .05$ , \*\* $p < .01$ , \*\*\* $p < .001$ . Degrees of freedom for the fixed effects were estimated using Satterthwaite's approximation.

#### Error-Related ICPS and Post-Error Attentional Control

Our LMM jointly testing whether error-related theta ICPS between midfrontal and the three lateral scalp regions predicted post-error attentional control (Table 4) revealed a significant main effect of error-related midfrontal-posterolateral ICPS. Specifically, increased error-related midfrontal-posterolateral ICPS was associated with higher post-error attentional control, as indexed by the reversed SSP-DDM attentional ratio (sd_a_/r_d_) (β = 0.17, 95% CI [0.05, 0.29], p = .007) (Figure 6). In contrast to our hypotheses, within the same statistical model, we found no significant effects involving midfrontal-frontolateral ICPS (all p >= .156; Table 4). Effects involving midfrontal-midlateral ICPS, condition, age, or their interactions were not significant (all p >= .085; Table 4).

**Figure 6.**
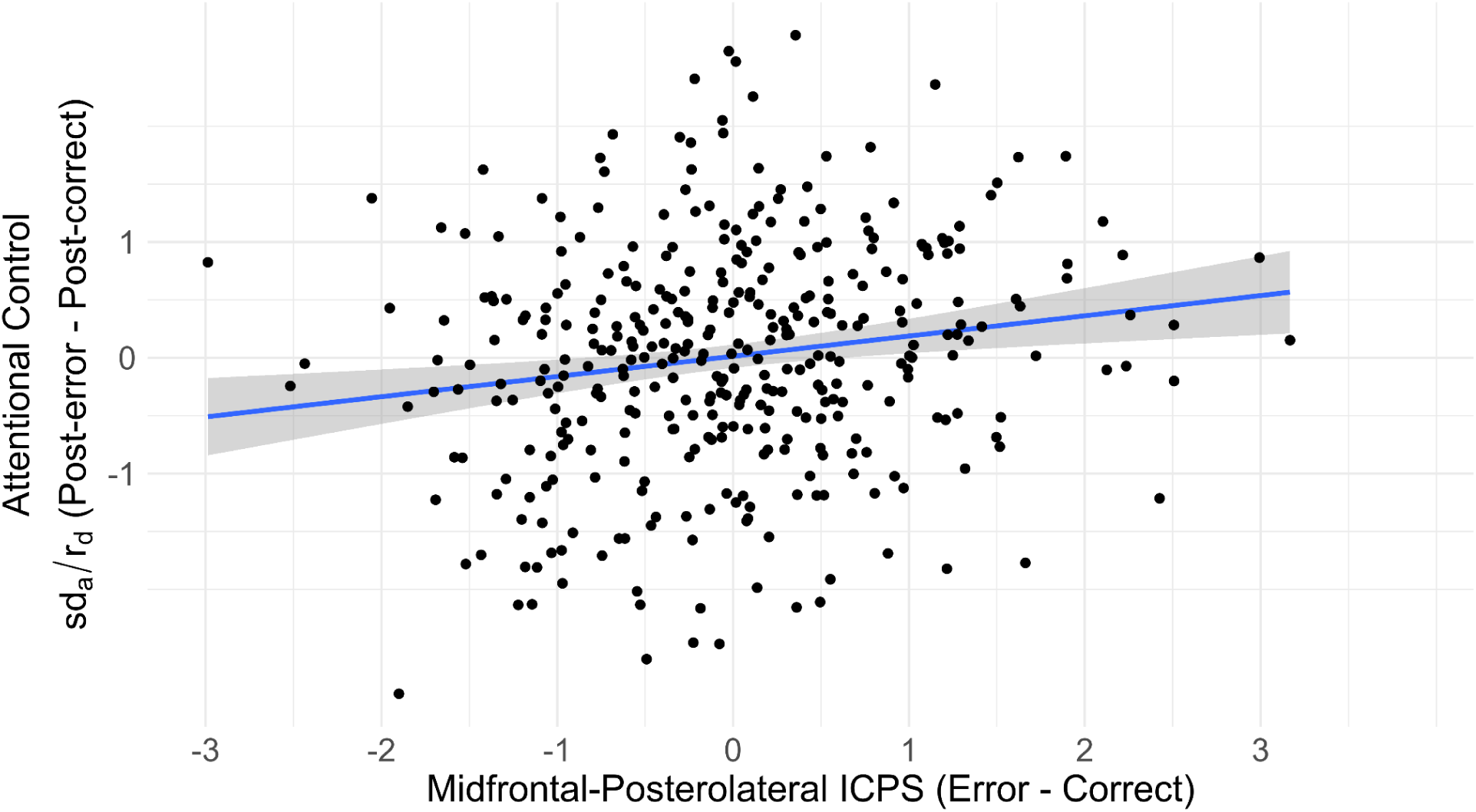
Relationship Between Error-Related Midfrontal-Posterolateral Theta ICPS and Reversed SSP-DDM Attentional Ratio (sd_a_/r_d_) *Note.* Shaded regions represent 95% confidence intervals.

#### Error-Related ICPS and Post-Error Response Caution

Contrary to our hypotheses, our LMM jointly testing whether error-related theta ICPS between midfrontal and the three lateral scalp regions predicted post-error response caution (boundary separation, a) revealed no significant main effects or interactions involving midfrontal-midlateral theta ICPS (all p >= .225; Table 5). Thus, our results provide no evidence that theta synchrony between midfrontal and midlateral regions increases response caution after errors. However, the model revealed a significant interaction between midfrontal-frontolateral ICPS and age (β = -0.15, 95% CI [-0.29, -0.01], p = .044) predicting post-error boundary separation (*a*; Figure 7). The nature of this interaction was such that error-related midfrontal-frontolateral theta ICPS exhibited a relatively more positive association with post-error boundary separation (response caution) at younger ages, whereas the opposite was true at relatively older ages. However, follow-up simple slope analyses (M age ± 1 SD) were not significant. Further effects involving midfrontal-posterolateral ICPS, condition, age, or their interactions were also not significant (all p >= .082; Table 5).

**Figure 7.**
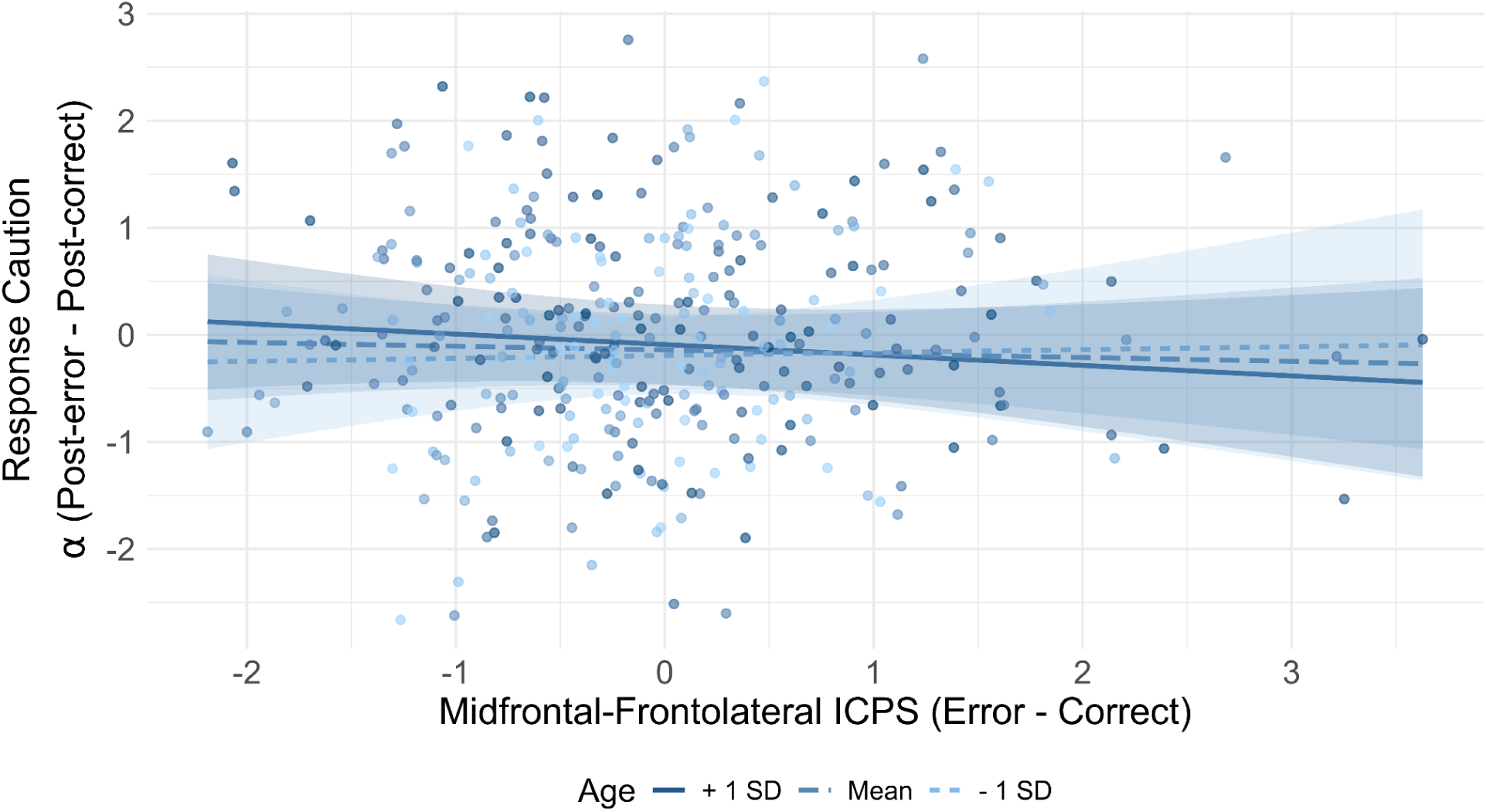
Interaction Between Error-Related Midfrontal-Frontolateral Theta ICPS and Age on Post-Error Response Caution (Boundary Separation, a) *Note.* Shaded regions represent 95% confidence intervals.

## Discussion

The current study investigated how error-related theta inter-regional synchrony between midfrontal and lateral scalp locations changes with age and social observation across early-mid adolescence. Further, we explored the functional significance of these inter-regional synchrony dynamics by testing relations with two forms of post-error control: attentional control and response caution, using measures derived from the SSP-DDM. We found a robust age-related increase in error-related theta inter-regional synchrony in two locations: midfrontal-frontolateral and midfrontal-midlateral; these effects were not modified by social observation. Additionally, we identified a significant interaction between age and social observation in predicting error-related midfrontal-posterolateral theta synchrony, such that error-related midfrontal-posterolateral theta synchrony significantly increased with age, but only within the non-social condition. Further, in our analyses of post-error adjustments, we found significant positive relationships between error-related midfrontal-posterolateral theta synchrony and our computational measure of post-error attentional control (*sd_a_/r_d_*). Additionally, we found that error-related midfrontal-frontolateral theta synchrony predicted post-error boundary separation (*a*), although this relation differed as a function of age. Taken together, these results extend existing findings on how the error monitoring system changes across adolescence, providing further insight into the role of social observation and the functional role of error-related theta inter-regional synchrony.

The first goal of this study was to close gaps in the literature regarding whether and how error-related theta inter-regional synchrony changes as a function of age and social observation during the adolescent period. Prior EEG work in youth, predominantly focusing on midfrontal activity, has consistently reported an age-related increase in error-related responses, measured both with event-related potentials (ERP) (Ladouceur et al., 2007; Tamnes et al., 2013) and TF measures (Morales et al., 2022; Morales & Buzzell, 2025). However, error-related midfrontal activity is thought to reflect only the first stage of error processing, reflecting error detection and signaling the need for additional control (Morales & Buzzell, 2025). Error-related theta inter-regional synchrony (indexed via ICPS) is thought to reflect the second stage of error processing, associated with the recruitment/instantiation of post-error control (Morales & Buzzell, 2025). Our findings demonstrate that error-related theta inter-regional synchrony significantly increased with age during early-to-mid adolescence (ages 10-14 years). To our knowledge, this is the first study to investigate age-related changes in error-related theta inter-regional synchrony (ICPS) across adolescence, as the few existing studies that examined such effects in developmental samples did not assess age-related changes (Buzzell et al., 2019) or did not capture the early-to-mid adolescent age range (Morales et al., 2022). Moreover, as we also identified relationships between error-related theta inter-regional synchrony and post-error adjustments (discussed below), our findings of age-related increases in error-related theta inter-regional synchrony provide insight into the possible development of post-error control instantiation networks beyond mere error detection. Below, we further discuss the role of social observation and the functional significance of error-related theta synchrony between midfrontal and lateral scalp locations.

Based on findings that adolescence is characterized by increased sensitivity to peer influence (Albert et al., 2013; Somerville, 2013; Steinberg & Monahan, 2007) and previous adolescent work reporting enhanced error-related brain responses under social observation (Barker et al., 2018; Buzzell et al., 2017, 2019; Hajcak et al., 2005; Kim et al., 2005; Voegler et al., 2018), we tested whether error-related theta inter-regional synchrony differed as a function of social observation. Among three spatial locations tested, social observation was found to selectively moderate age-related increases in error-related midfrontal-posterolateral theta synchrony. Specifically, we only observed age-related increases in error-related midfrontal-posterolateral theta synchrony within the non-social condition. One tentative hypothesis is that the lack of age-related increases in error-related responses in the social condition may reflect social observation being a source of additional cognitive load that limits the capacity to appropriately engage error-related theta inter-regional synchrony responses. Such moderation of the age effect on error-related theta synchrony by social observation might reflect increasing sensitivity to social evaluation throughout adolescence (Somerville, 2013) making the social condition potentially more taxing for older adolescents. For example, fMRI studies have shown that adolescents in social context, compared to adults, exhibit greater activity in regions associated with mentalizing, including the medial prefrontal cortex (mPFC) (Blakemore et al., 2007; Burnett et al., 2009; Pfeifer et al., 2009; Somerville et al., 2013) and the temporoparietal junction (TPJ) (Pfeifer et al., 2009), as well as enhanced connectivity between these regions (Burnett et al., 2009). Furthermore, increased activation in the TPJ while under social observation has been shown to negatively correlate with activity in error monitoring regions such as the dorsal anterior cingulate cortex (dACC) (Peake et al., 2013). Moreover, social observation in adolescence induces greater activity in affect-regulation regions (Ahmed et al., 2015; Andrews et al., 2021), including the ventral striatum (VS) and orbitofrontal cortex (OFC) (Chein et al., 2011), suggesting a link between social evaluation and other emotionally salient contexts. Importantly, the processing of emotional cues has been shown to induce greater activity in frontostriatal connections (Somerville et al., 2011) and to impact performance on cognitive control tasks (Ahmed et al., 2015; Somerville et al., 2011). Taken together, these neural interactions occurring under social observation may increase cognitive load and interfere with overlapping neural activity that is indexed by midfrontal-posterolateral theta synchrony, limiting the overall capacity for error monitoring and post-error control.

Having characterized the effects of age and social observation on distinct forms of error-related theta inter-regional synchrony, we next sought to determine the functional significance of each of these measures by testing relations with SSP-DDM measures of post-error attentional control and response caution. We found that increased midfrontal-posterolateral theta synchrony was associated with an increase in post-error selective attention (*sd_a_/r_d_*), regardless of age or social observation. These findings are consistent with empirical data in adults from Cohen and colleagues (2009), who showed that error-related theta synchrony between midfrontal and occipital regions following errors predicts raw behavioral measures that are thought to similarly reflect attentional control (ratio of raw RT and accuracy) in post-error trials. Our findings are also in line with the broader notion that the medial frontal cortex is involved in attention networks that propagate top-down attentional control from midfrontal to occipitoparietal regions, as demonstrated by EEG studies examining cross-frequency coupling between midfrontal theta and occipitoparietal gamma (Moser et al., 2024) and alpha (Cohen & van Gaal, 2013). Similarly, in fMRI studies employing the Stroop task, processing stimulus-response conflict has been associated with increased functional connectivity between the ACC and parietal regions (Mayer et al., 2012), and crucially, such connectivity increases have been linked to improved task accuracy (Rosenberg-Katz et al., 2016). However, to our knowledge, the current study is the first to demonstrate the role of error-related theta synchrony between midfrontal and posterolateral regions in post-error attentional control using the SSP-DDM.

We also hypothesized that error-related midfrontal-frontolateral theta synchrony would relate to post-error attentional control. Prior work has linked error-related theta synchrony between midfrontal and frontolateral regions to raw behavioral measures of post-error adjustments, such as PES (Cavanagh et al., 2009) and PERI (Buzzell et al., 2019). Contrary to our hypotheses, we found no effects of error-related midfrontal-frontolateral theta synchrony on post-error attentional control (*sd_a_/r_d_*). Instead, we only found non-specific relations between error-related midfrontal-frontolateral theta synchrony and age on post-error response caution (*a*). That is, error-related midfrontal-frontolateral theta synchrony demonstrated a relatively more positive relationship with post-error response caution in younger adolescents, whereas the opposite trend was observed in older adolescents. One tentative hypothesis is that these changing associations between error-related midfrontal-frontolateral theta synchrony and post-error adjustments may occur due to ongoing development of the prefrontal cortex and remodelling of its connections with other brain regions (Ahmed et al., 2015; Luna et al., 2015). In this view, the interaction of error-related midfrontal-frontolateral theta synchrony with age may be a result of developmental shifts in the functional organization of midfrontal-prefrontal connections, such that adolescents of different ages may recruit differential mechanisms of cognitive control through similar neural regions. Although speculative, this interpretation is consistent with longitudinal fMRI studies that show developmental shifts in prefrontal and medial frontal activation patterns across adolescence, which correspond to changes in behavioral strategies in cognitive control tasks (Kim-Spoon et al., 2021; Luna et al., 2013; McCormick et al., 2017). However, future longitudinal research is required to determine whether the opposing behavioral shifts we observed in this cross-sectional sample represent true developmental trajectories associated with cognitive control.

Given previous findings demonstrating that error-related midfrontal-midlateral theta synchrony is associated with increased PES (Buzzell et al., 2019), we hypothesized its role in post-error response caution. However, contrary to our hypotheses, we found no relationship between error-related midfrontal-midlateral theta synchrony and post-error response caution (*a*) indexed by SSP-DDM. Nonetheless, in supplemental analyses we replicated the link between midfrontal-midlateral theta synchrony and the raw PES behavioral measure (see Supplemental Materials). The finding that midfrontal-midlateral theta synchrony was not associated with response caution is consistent with the broader notion that PES emerges at least partially from a reactive orienting response (Notebaert et al., 2009), which may already be functionally mature in adolescence (Valadez et al., 2022). In contrast, the strategic adjustment of decision boundaries relies on more strategic control mechanisms that may undergo a more protracted developmental trajectory and continue to exhibit large variability throughout adolescence (Fosco et al., 2022). This discrepancy further highlights that raw behavioral measures of post-error adjustments (e.g., PES) are not specific regarding the cognitive processes they actually reflect (Draheim et al., 2019; Dutilh et al., 2019; White & Kitchen, 2022).

Of note, the analyses of error-related theta inter-regional synchrony and its relationships with post-error adjustments exhibited only limited effects of social observation. One possibility is that the limited effects of social observation were a result of the specific social observation paradigm employed in the current study. As noted by Stibolt and colleagues (2026), prior work suggests that the effects of social observation on error monitoring may depend on factors such as the observer’s location (physical/virtual presence and/or physical proximity; e.g., Barker et al., 2015; Schillinger et al., 2016; Voegler et al., 2018), the observer’s behavior (e.g., whether they marked every mistake or simply remained present; Barker et al., 2015; Voegler et al., 2018), and the observer’s identity (peer/authority figure; e.g., Claypoole & Szalma, 2017). For the current study, adolescent participants were observed by a similarly-aged peer located in a separate room and via video call. Peer observers were limited in their ability to evaluate the correctness of individual responses (trials) and read aloud the block-level feedback. This approach was employed to limit interactions between the participants during non-social portions of the study and to standardize the social condition experience across participants/sessions. Future work should seek to further explore how these additional factors might impact the effects of social observation on error-related theta inter-regional synchrony.

## Limitations and Future Directions

Given that the current study focused on an early-to-mid adolescent age range (10-14 years), it remains unknown whether the observed effects will generalize across development. Nevertheless, our findings on error-related theta dynamics and their functional role in post-error control align closely with data from young adult populations (Cavanagh et al., 2009; Cohen et al., 2009; Cohen & van Gaal, 2013; Driel et al., 2012). Additionally, this study utilized a cross-sectional design, which limits our ability to draw strong inferences regarding developmental phenomena. Thus, future work should utilize longitudinal data across a broader age range to fully capture these developmental trajectories over time.

## Conclusion

In this report, we investigated age-related and social observation effects on error-related theta dynamics, as well as explored their functional role in adolescent post-error behavior. We first characterized age-related changes in error-related theta synchrony, finding that error-related midfrontal-frontolateral and midfrontal-midlateral theta synchrony significantly increased with age. We also showed that age-related changes in error-related midfrontal-posterolateral theta synchrony, in particular, are modulated by social observation. Using computational modeling, we demonstrated that error-related midfrontal-posterolateral theta synchrony is predictive of post-error attentional control regardless of age and social observation manipulations, whereas error-related midfrontal-frontolateral theta synchrony relates to post-error response caution—albeit dependent on age. Collectively, our findings indicate dynamic maturational and social observation changes in error-related theta inter-regional synchrony responses and provide evidence for their role in behavioral adaptation across adolescence.

## Data and Code Availability

The scripts for data pre- and post-processing, as well as the statistical analysis scripts, are available on the following GitHub repository: https://github.com/NDCLab/thrive-theta-ddm.

## Declaration of Generative AI

During the preparation of this work, the authors used Google Gemini 2.5 Pro for limited aspects of computer programming and to refine the clarity of certain passages. After using these tools/services, the authors reviewed and edited the content as needed and take full responsibility for the content of the published article.

## Acknowledgement

We are deeply grateful to all participants and their families for their time, engagement, and willingness to contribute to this research.

## Author Note

Research reported in this publication was supported by the National Institute of Mental Health of the National Institutes of Health under award number R01MH131637 (Buzzell, Pettit). FZ is grateful for the support he received through the FIU UGS Presidential Fellowship during his PhD study.

