## Supplementary Materials for "Age and Social Observation Effects on Theta Synchrony and Its Role in Adolescent Post-Error Control: A Computational Approach"

### In-Person Visit Social Interaction Protocol

For sessions where both participants were in-person, the dyad partners performed all tasks in separate rooms, and all instructions and study procedures were delivered by different research assistants (RAs). During active task blocks across all conditions, the RA was outside the testing room with the door closed to prevent unintended social presence or distraction. The RA continuously monitored the participant's behavioral responses and EEG data. The RA only re-entered the testing room between task blocks to correct EEG artifacts (e.g., excessive blinking or muscle tension, limited to three reminders, or an increase in impedance in a selected electrode).

Prior to the start of the social condition, the RA assigned to the performing participant delivered a standardized introduction between the dyad partners over the Zoom call while being in the room with the performing participant. To verify visual clarity, the observer was required to read a word displayed on the performing participant's stimulus screen. Once visibility and audio fidelity were confirmed, standardized instructions for the Flanker task were delivered to the performing participant, which included performing the task as quickly and as accurately as possible and a set of additional instructions to minimize EEG artifacts, such as limiting eye blinks and avoiding excessive muscle activity. The observer was instructed to closely monitor their dyad partner's performance and to verbally read the performance feedback presented on the screen aloud after each block of trials. The observer's microphone remained on to facilitate this feedback, but the observer was explicitly directed to avoid making any other distracting noises during the task. The performing participant was instructed to verbally acknowledge the receipt of this feedback after each block by confirming they heard it, or by stating if no feedback was heard. If the observer failed to read the feedback aloud, the RA assigned to the observer participant intervened between blocks by entering the room, temporarily muting the microphone, and providing a targeted reminder.

During the social condition, the iPad with the Zoom video call was attached to a floor-mounted stand positioned behind and to one side of the performing participant's chair. To ensure a standardized viewing angle for the observer, the camera was angled specifically to capture the performing participant's hands resting on the button box, as well as the stimulus computer screen. Conversely, for the observer, the iPad was located on a desk stand placed directly in front of them, ensuring the observer's head and shoulders remained centered in the frame. For both participants, self-view and visual tracking features, including "Center Stage" and "Portrait" effects on the iPad's camera, were disabled. Outside the social parts of the study, the Zoom iPad's camera and audio were disabled, and the iPad was physically covered with a black sheet to establish a strict non-social baseline and to ensure participant privacy.

**Details on EEG Preprocessing**

EEG data was preprocessed in MATLAB R2021b (MathWorks Inc., Sherborn, MA, USA) using the EEGLAB toolbox (Delorme & Makeig, 2004) and a modified MADE pipeline (Debnath et al., 2020). Based on timing synchronization tests, stimulus offsets were corrected by 12 ms, while response offsets (measured at < 1 ms) required no adjustment. The continuous data were high-pass filtered at 0.1 Hz, followed by a low-pass of 49 Hz and a stopband of 59 Hz. Bad channels were identified and excluded using the FASTER plugin (Nolan et al., 2010). To isolate and remove ocular and/or muscular artifacts, we employed independent component analysis (ICA). To optimize the ICA decomposition, we followed established protocols (Debnath et al., 2020) by creating a temporary copy of each dataset, high-pass filtering it at 1 Hz, and segmenting it into 1-second epochs. Channels containing artifacts in more than 20% of these epochs were removed from both the copied and original datasets. Following the ICA on the filtered/cleaned copy, the resulting weights were transferred back to the original dataset. Artifactual independent components (ICs) were then identified using the adjusted-ADJUST algorithm (Leach et al., 2020; Mognon et al., 2011) and subtracted from the data. After the ICA, the data were segmented into 3-second epochs (-1000 to 2000 ms relative to the response), and the DC offset was subtracted from each epoch. Remaining artifacts were addressed using a ±125 µV threshold: epochs were rejected if this threshold was exceeded in channels near the eyes. For all other channels not located near the eyes, values exceeding this threshold were interpolated, unless more than 10% of the channels in a single epoch reached the limit, in which case the entire epoch was discarded. Finally, missing channels were reconstructed via spherical spline interpolation, and all data were re-referenced to the common average.

**SSP-DDM Fitting and Parameter Recovery Simulations**

SSP-DDM fitting was conducted using C++ code executed in R (R Core Team, 2021) via the Rcpp package (Eddelbuettel & Francois, 2011) using modified code originally written for the flankr package (Grange, 2016). The model was fit separately for each participant, for each trial type (post-error and post-correct), and each condition (social/non-social). The upper and lower bounds of SSP-DDM parameter values used for fitting were taken from a similar SSP-DDM parameter recovery study (White et al., 2018). Each fitting iteration included a simulation of 10000 trials for each trial type (congruent/incongruent) using a set of model parameters; comparison with the actual data (RT distributions) involved $\chi^2$ calculation that simultaneously accounted for congruent/incongruent data. Minimization of the $\chi^2$ value was performed via a differential evolution optimization algorithm using the DEoptim R package (Mullen et al., 2011).

To determine a minimum sufficient number of post-error trials for subjects to be included in the DDM-related analyses, a parameter recovery simulation analysis was conducted prior to fitting the real data. For each number of post-error trials (N = 12, 16, 20, 24), 300 participants were simulated, and the model was fit using the same routine described above, and then the recovered parameters were correlated with the original parameters using Spearman correlation coefficient (Figure S1). Correlation values >= .6 between original and recovered parameters were considered acceptable.

**Figure S1**

*SSP-DDM Recovery Results From 300 Simulations*

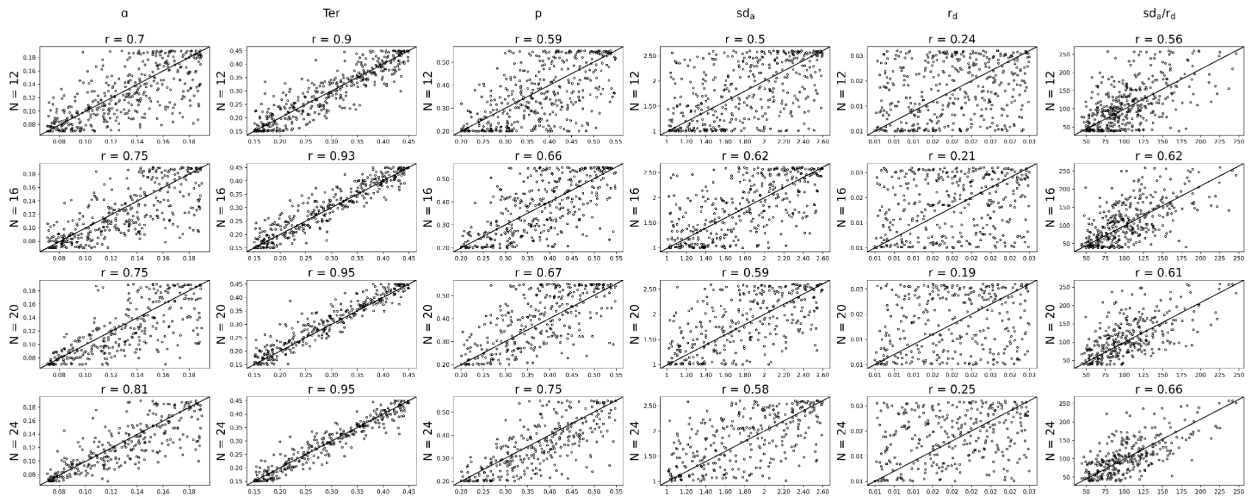


*Note.* Simulated values are plotted against recovered values for each parameter for a different number of post-error trials (N). The quality of recovery was quantified using Spearman correlation coefficient (r).

### Preliminary Analyses of Behavior and EEG

The primary focus of this study was to investigate the effects of age and social observation on error-related theta inter-regional synchrony (ICPS) and its relations with post-error adjustments. Nonetheless, we first performed a series of preliminary analyses to ensure expected patterns of raw flanker task behavior (proportion correct and RT on the current trial) and error-related theta responses (power and ITPS) over the midfrontal region, in line with prior work (see (Eriksen & Eriksen, 1974) for behavior and (Buzzell et al., 2019; Cavanagh et al., 2009; Cavanagh & Frank, 2014) for EEG).

#### *Flanker Task Behavior*

Prior to performing statistical analyses on raw flanker task behavioral measures, data were preprocessed using a similar approach as used prior to DDM fitting. Additionally, only correct trial data were included in the RT statistical model to isolate the effect of congruency and eliminate its confound with accuracy (Buzzell et al., 2019). All analyses involving RT were conducted on log-transformed RT values to adjust for the positive skew of RT distributions (Miller, 1988). Thus, both models included binary variables for congruency (congruent/incongruent) and condition (social/non-social), and a continuous variable for age in months as fixed effects with interactions, differing only in their outcome variables (mean proportion correct/mean RT). See Table S1 below for numbers of trials and participants included in the analyses.

**Table S1**

*Number of Trials and Participants Included in the Analyses of Flanker Task Behavior*

| Data type (measure) | NS Incon. | NS Con. | S Incon. | S Con. |
|---|---|---|---|---|
| Trials per Participant (M ± SD) | | | | |
| Behavior | 149.12 ± 19.01 | 184.18 ± 11.34 | 149.46 ± 17.79 | 184.89 ± 11.29 |
| Participants per Analysis | | | | |
| Proportion Correct | 223 | 222 | 227 | 224 |
| RT | 222 | 220 | 226 | 225 |

*Note.* "S" denotes social condition, and "NS" denotes nonsocial condition.

Consistent with congruency effects expected in a flanker task (Eriksen & Eriksen, 1974), there was a significant main effect of congruency on proportion correct (β = 0.77, 95% CI [0.73, 0.80], $p < .001$; Table S1) and RT (β = -0.43, 95% CI [-0.45, -0.40], $p < .001$; Table S2), such that the proportion correct was higher and RTs were faster in congruent trials. Additionally, there was a significant main effect of age on RT (β = -0.34, 95% CI [-0.44, -0.24], $p < .001$), with increasing age corresponding to faster RT. Furthermore, there was a significant age * congruency interaction (β = 0.04, 95% CI [0.01, 0.06], $p = .003$), such that the decrease in RT with age was significantly greater for incongruent (β = -.38, 95% CI [-0.50, -0.26], $p < .001$) compared to congruent (β = -.30, 95% CI [-0.42, -0.18], $p < .001$) trials.

**Table S2**

*Flanker Task Proportion Correct Statistics*

| Parameter | *β* | *SE* | *df* | *t* | *p* | 95% CI |
|---|---|---|---|---|---|---|
| (Intercept) | 0.03 | 0.04 | 229.64 | 0.63 | .529 | [-0.05, 0.11] |
| Congruency | 0.77 | 0.02 | 647.43 | 42.78 | < .001*** | [0.73, 0.80] |
| Condition | 0.02 | 0.02 | 684.52 | 0.89 | .375 | [-0.02, 0.05] |
| Age | -0.01 | 0.03 | 229.38 | -0.25 | .801 | [-0.06, 0.05] |
| Sex | 0.05 | 0.03 | 227.77 | 1.76 | .080 | [-0.01, 0.11] |
| Peer mode | -0.04 | 0.04 | 228.86 | -1.00 | .317 | [-0.12, 0.04] |
| Congruency * Condition | 0.01 | 0.02 | 647.45 | 0.29 | .771 | [-0.03, 0.04] |
| Congruency * Age | 0.03 | 0.02 | 647.71 | 1.76 | .078 | [-0.00, 0.07] |

| | | | | | | |
|---|---|---|---|---|---|---|
| Condition * Age | 0.01 | 0.02 | 688.19 | 0.53 | .598 | [-0.03, 0.05] |
| Congruency * Condition * Age | -0.01 | 0.02 | 647.71 | -0.68 | .500 | [-0.05, 0.02] |

*Note.* Significance codes: * p < .05, ** p < .01, *** p < .001. Degrees of freedom for the fixed effects were estimated using Satterthwaite's approximation.

**Table S3**

*Flanker Task RT Statistics*

| Parameter | *β* | *SE* | *df* | *t* | *p* | 95% CI |
|---|---|---|---|---|---|---|
| (Intercept) | 0.06 | 0.07 | 231.18 | 0.86 | .390 | [-0.08, 0.21] |
| Congruency | -0.43 | 0.01 | 644.15 | -33.49 | < .001*** | [-0.45, -0.40] |
| Condition | -0.02 | 0.01 | 653.68 | -1.39 | .165 | [-0.04, 0.01] |
| Age | -0.34 | 0.05 | 230.50 | -6.65 | < .001*** | [-0.44, -0.24] |
| Sex | 0.13 | 0.05 | 230.44 | 2.53 | .012* | [0.03, 0.23] |
| Peer mode | -0.05 | 0.07 | 230.91 | -0.68 | .497 | [-0.19, 0.09] |
| Congruency * Condition | -0.00 | 0.01 | 644.15 | -0.31 | .754 | [-0.03, 0.02] |
| Congruency * Age | 0.04 | 0.01 | 643.97 | 2.93 | .003** | [0.01, 0.06] |
| Condition * Age | -0.02 | 0.01 | 653.86 | -1.43 | .153 | [-0.04, 0.01] |
| Congruency * Condition * Age | 0.00 | 0.01 | 643.97 | 0.14 | .889 | [-0.02, 0.03] |

*Note.* Significance codes: * p < .05, ** p < .01, *** p < .001. Degrees of freedom for the fixed effects were estimated using Satterthwaite's approximation.

#### ***Midfrontal Theta Power and ITPS***

As preliminary measures, response-locked midfrontal theta power and intertrial phase synchrony (ITPS) were computed in the midfrontal cluster of electrodes (FCz and surrounding electrodes: 1, 2, 33, 34 on Figure 2). For midfrontal theta power, TF power was computed for each epoch for each channel, and then epochs were averaged. ITPS was computed as the average of phase angle differences between trials in the midfrontal electrode cluster and could vary from 0 to 1 (with 0 corresponding to a uniform distribution of phase angles, and 1 denoting perfect phase alignment). To compute ITPS, a subsampling algorithm identical to the one used for ICPS was employed. After TF decomposition, all subsequent processing as well as inclusion criteria for participants, individual epochs, and statistical analyses were identical to ICPS analyses reported in the main text.

For the theta power model, the outcome variable was mean midfrontal theta power. Binary variables for accuracy (correct/incorrect) and condition (social/non-social), and a continuous variable for age in months were included as fixed effects with interactions. The theta ITPS model used identical predictors but with mean midfrontal theta ITPS as the outcome. Below, we report results on previously established effects of accuracy (Buzzell et al., 2019; Cavanagh et al., 2009; Cavanagh & Frank, 2014), social observation (Buzzell et al., 2019), and age effects for these measures.

***Theta Power Results***

The LMM (Table S3) revealed a significant main effect of accuracy (β = -0.74, 95% CI [-0.77, -0.70], $p < .001$) greater power in response to errors compared to corrects (Figure S2), a positive main effect of age (β = 0.17, 95% CI [0.11, 0.23], $p < .001$), and age * accuracy interaction (β = -0.09, 95% CI [-0.12, -0.05], $p < .001$). Follow-up analysis showed that the age effect was significantly greater in error-related (β = 0.25, 95% CI [0.18, 0.33], $p < .001$) than in correct-related (β = 0.08, 95% CI [0.01, 0.15], $p = .022$) responses. The effect of the condition was not significant (β = 0.01, 95% CI [-0.02, 0.05], $p = .448$).

**Table S4**

*Midfrontal Theta Power Statistics*

| Parameter | *β* | *SE* | *df* | *t* | *p* | 95% CI |
|---|---|---|---|---|---|---|
| (Intercept) | 0.02 | 0.04 | 226.91 | 0.34 | .735 | [-0.07, 0.10] |
| Accuracy | -0.74 | 0.02 | 623.34 | -41.75 | < .001*** | [-0.77, -0.70] |
| Condition | 0.01 | 0.02 | 653.12 | 0.76 | .448 | [-0.02, 0.05] |
| Age | 0.17 | 0.03 | 228.83 | 5.42 | < .001*** | [0.11, 0.23] |
| Sex | -0.12 | 0.03 | 226.57 | -3.89 | < .001*** | [-0.18, -0.06] |
| Peer mode | -0.00 | 0.04 | 226.15 | -0.09 | .932 | [-0.09, 0.08] |
| Accuracy * Condition | -0.01 | 0.02 | 622.29 | -0.62 | .535 | [-0.05, 0.02] |
| Accuracy * Age | -0.09 | 0.02 | 623.47 | -4.90 | < .001*** | [-0.12, -0.05] |
| Condition * Age | 0.00 | 0.02 | 657.89 | 0.13 | .897 | [-0.03, 0.04] |
| Accuracy * Condition * Age | 0.02 | 0.02 | 622.66 | 1.24 | .214 | [-0.01, 0.06] |

*Note.* Significance codes: * p < .05, ** p < .01, *** p < .001. Degrees of freedom for the fixed effects were estimated using Satterthwaite's approximation.

**Figure S2**

*Response-Locked Midfrontal Theta Power in Two Conditions*

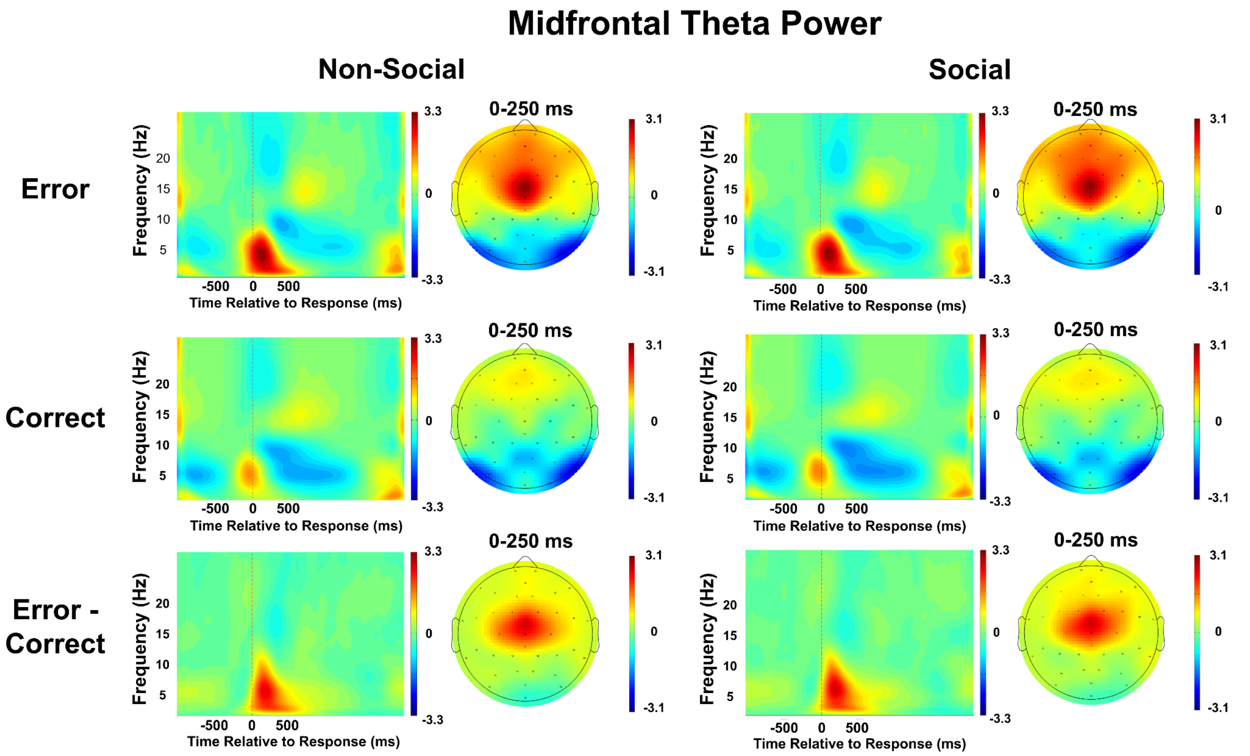


***Theta ITPS Results***

The LMM (Table S4) revealed a significant main effect of accuracy (β = -0.18, 95% CI [-0.24, -0.13], $p < .001$) with greater ITPS in response to errors compared to corrects (Figure S3), a positive main effect of age (β = 0.14, 95% CI [0.06, 0.23], $p$ = 001), and age * accuracy interaction (β = -0.11, 95% CI [-0.17, -0.05], $p < .001$). Follow-up analysis showed that the age effect was only significant for error-related (β = 0.25, 95% CI [0.15, 0.36], $p < .001$) but not for correct-related (β = 0.04, 95% CI [-0.07, 0.14], $p = .482$) responses. Additionally, there was a significant effect of biological sex such that the ITPS was greater in females (β = 0.09, 95% CI [0.00, 0.17], $p = .045$). The effect of the condition was not significant (β = 0.00, 95% CI [-0.05, 0.06], $p = .879$).

**Table S5**

*Midfrontal Theta ITPS Statistics*

| Parameter | *β* | *SE* | *df* | *t* | *p* | 95% CI |
|---|---|---|---|---|---|---|

| | | | | | | |
|---|---|---|---|---|---|---|
| (Intercept) | -0.01 | 0.06 | 221.66 | -0.11 | .913 | [-0.13, 0.11] |
| Accuracy | -0.18 | 0.03 | 617.82 | -6.31 | < .001*** | [-0.24, -0.13] |
| Condition | 0.00 | 0.03 | 655.16 | 0.15 | .879 | [-0.05, 0.06] |
| Age | 0.14 | 0.04 | 223.28 | 3.41 | .001*** | [0.06, 0.23] |
| Sex | 0.09 | 0.04 | 220.38 | 2.02 | .045* | [0.00, 0.17] |
| Peer mode | 0.00 | 0.06 | 220.80 | 0.06 | .952 | [-0.12, 0.12] |
| Accuracy * Condition | 0.02 | 0.03 | 616.52 | 0.68 | .500 | [-0.04, 0.08] |
| Accuracy * Age | -0.11 | 0.03 | 617.73 | -3.71 | < .001*** | [-0.17, -0.05] |
| Condition * Age | -0.02 | 0.03 | 662.40 | -0.51 | .610 | [-0.07, 0.04] |
| Accuracy * Condition * Age | 0.02 | 0.03 | 616.67 | 0.52 | .605 | [-0.04, 0.07] |

*Note.* Significance codes: * p < .05, ** p < .01, *** p < .001. Degrees of freedom for the fixed effects were estimated using Satterthwaite's approximation.

**Figure S3**

*Response-Locked Midfrontal Theta ITPS in Two Conditions*

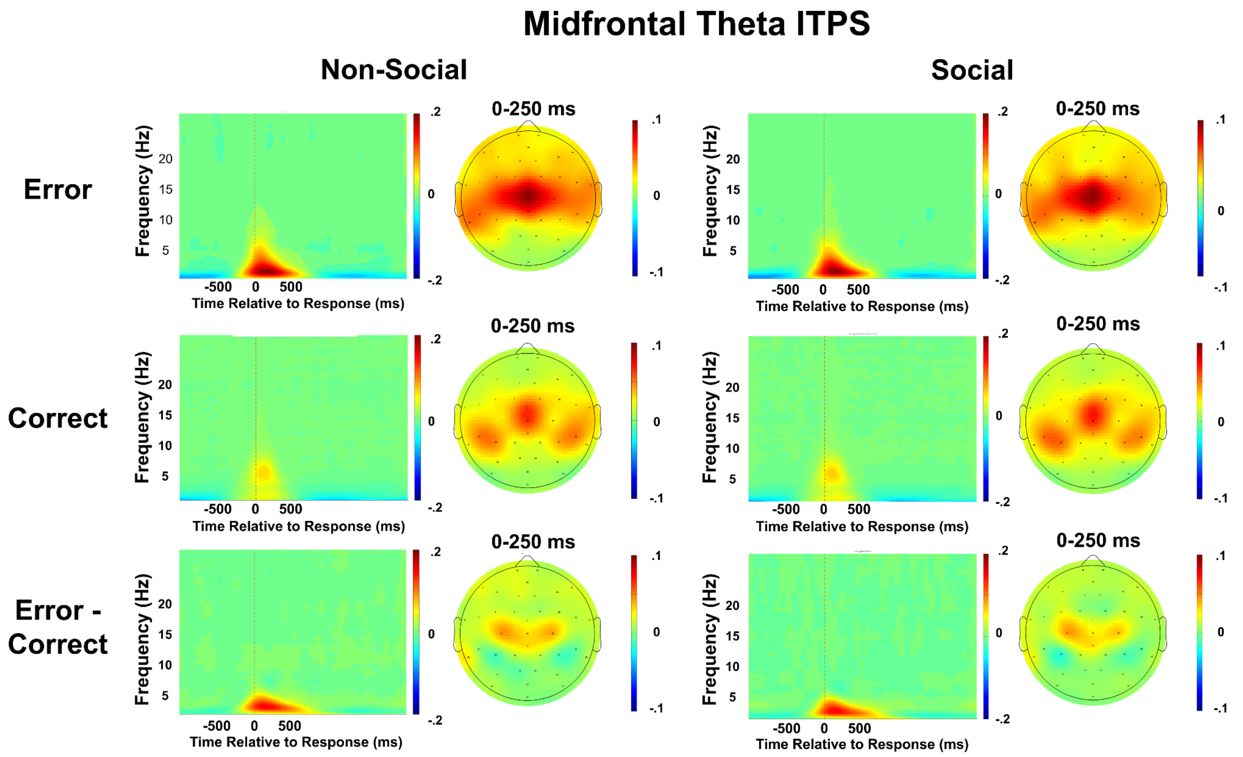


**Exploratory Analyses on Functional Role of Error-Related Theta ICPS in Post-Error Control**

In the main text, we reported the results of two models, each simultaneously testing the effects of three error-related theta ICPS measures (midfrontal-frontolateral, midfrontal-midlateral, and midfrontal-posterolateral) on post-error attentional control ($sd_a/r_d$) or response caution (*a*). Below, we provide the full outputs of the six more exhaustive exploratory models in which we predicted each of the two SSP-DDM measures ($sd_a/r_d$ and $a$) using each of the three theta ICPS measures entered as predictors within separate models (Tables S5-S10). These six exploratory models revealed no qualitative differences compared to the full models reported in the main text.

**Table S5**

*Midfrontal-Frontolateral ICPS Predicting SSP-DDM Attentional Ratio ($sd_a/r_d$) Model Statistics*

| Parameter | *β* | *SE* | *df* | *t* | *p* | 95% CI |
|---|---|---|---|---|---|---|
| (Intercept) | -0.04 | 0.08 | 212.67 | -0.51 | .609 | [-0.21, 0.12] |
| ICPS frontolateral | 0.04 | 0.06 | 347.37 | 0.72 | .474 | [-0.07, 0.16] |
| Condition | -0.07 | 0.05 | 179.60 | -1.31 | .191 | [-0.16, 0.03] |
| Age | -0.01 | 0.06 | 193.69 | -0.11 | .916 | [-0.12, 0.11] |
| Sex | -0.04 | 0.06 | 187.74 | -0.76 | .447 | [-0.15, 0.07] |
| Peer mode | 0.05 | 0.08 | 211.61 | 0.63 | .530 | [-0.11, 0.22] |
| ICPS frontolateral * Condition | -0.05 | 0.06 | 310.19 | -0.79 | .429 | [-0.16, 0.07] |
| ICPS frontolateral * Age | 0.02 | 0.06 | 349.59 | 0.27 | .787 | [-0.11, 0.14] |
| Condition * Age | 0.01 | 0.05 | 182.94 | 0.26 | .792 | [-0.09, 0.11] |
| ICPS frontolateral * Condition * Age | 0.04 | 0.06 | 331.09 | 0.59 | .552 | [-0.09, 0.16] |

*Note.* Significance codes: * p < .05, ** p < .01, *** p < .001. Degrees of freedom for the fixed effects were estimated using Satterthwaite's approximation.

**Table S6**

*Midfrontal-Midlateral ICPS Predicting SSP-DDM Attentional Ratio ($sd_a/r_d$) Model Statistics*

| Parameter | *β* | *SE* | *df* | *t* | *p* | 95% CI |
|---|---|---|---|---|---|---|
| (Intercept) | -0.06 | 0.08 | 208.46 | -0.72 | .474 | [-0.22, 0.10] |
| ICPS midlateral | 0.04 | 0.06 | 317.88 | 0.70 | .484 | [-0.07, 0.16] |
| Condition | -0.06 | 0.05 | 185.89 | -1.15 | .252 | [-0.16, 0.04] |
| Age | -0.03 | 0.06 | 198.26 | -0.45 | .656 | [-0.14, 0.09] |
| Sex | -0.03 | 0.06 | 193.26 | -0.57 | .570 | [-0.14, 0.08] |
| Peer mode | 0.06 | 0.08 | 205.65 | 0.69 | .491 | [-0.10, 0.21] |
| ICPS midlateral * Condition | 0.05 | 0.06 | 264.67 | 0.97 | .333 | [-0.05, 0.16] |
| ICPS midlateral * Age | 0.08 | 0.06 | 331.81 | 1.42 | .157 | [-0.03, 0.20] |
| Condition * Age | -0.00 | 0.05 | 190.10 | -0.01 | .989 | [-0.10, 0.10] |
| ICPS midlateral * Condition * Age | -0.00 | 0.06 | 289.35 | -0.01 | .994 | [-0.11, 0.11] |

*Note.* Significance codes: * p < .05, ** p < .01, *** p < .001. Degrees of freedom for the fixed effects were estimated using Satterthwaite's approximation.

**Table S7**

*Midfrontal-Posterolateral ICPS Predicting SSP-DDM Attentional Ratio ($sd_a/r_d$) Model Statistics*

| Parameter | *β* | *SE* | *df* | *t* | *p* | 95% CI |
|---|---|---|---|---|---|---|
| (Intercept) | -0.03 | 0.08 | 194.17 | -0.39 | .701 | [-0.18, 0.12] |
| ICPS posterolateral | 0.17 | 0.06 | 326.08 | 2.95 | .003** | [0.06, 0.28] |
| Condition | -0.05 | 0.05 | 176.64 | -1.01 | .313 | [-0.14, 0.05] |
| Age | 0.02 | 0.05 | 183.88 | 0.31 | .757 | [-0.09, 0.12] |
| Sex | -0.03 | 0.05 | 184.91 | -0.50 | .616 | [-0.13, 0.08] |
| Peer mode | 0.06 | 0.08 | 194.54 | 0.72 | .471 | [-0.09, 0.21] |
| ICPS posterolateral * Condition | 0.06 | 0.05 | 277.20 | 1.12 | .262 | [-0.04, 0.16] |
| ICPS posterolateral * Age | 0.03 | 0.06 | 345.83 | 0.52 | .604 | [-0.08, 0.14] |
| Condition * Age | 0.03 | 0.05 | 176.68 | 0.55 | .586 | [-0.07, 0.12] |
| ICPS posterolateral * Condition * Age | -0.01 | 0.06 | 303.09 | -0.14 | .891 | [-0.12, 0.10] |

*Note.* Significance codes: * p < .05, ** p < .01, *** p < .001. Degrees of freedom for the fixed effects were estimated using Satterthwaite's approximation.

**Table S8**

*Midfrontal-Frontolateral ICPS Predicting SSP-DDM Boundary Separation (a) Model Statistics*

| Parameter | *β* | *SE* | *df* | *t* | *p* | 95% CI |
|---|---|---|---|---|---|---|
| (Intercept) | -0.00 | 0.08 | 210.73 | -0.06 | .950 | [-0.16, 0.15] |
| ICPS frontolateral | 0.06 | 0.06 | 335.82 | 0.94 | .346 | [-0.06, 0.17] |
| Condition | -0.01 | 0.05 | 183.19 | -0.15 | .882 | [-0.11, 0.09] |
| Age | 0.10 | 0.05 | 186.87 | 1.83 | .068 | [-0.01, 0.21] |
| Sex | 0.11 | 0.05 | 183.49 | 2.14 | .033* | [0.01, 0.22] |
| Peer mode | 0.02 | 0.08 | 211.13 | 0.30 | .765 | [-0.13, 0.18] |
| ICPS frontolateral * Condition | 0.02 | 0.06 | 327.30 | 0.27 | .791 | [-0.10, 0.13] |
| ICPS frontolateral * Age | -0.18 | 0.06 | 344.84 | -2.90 | .004** | [-0.30, -0.06] |
| Condition * Age | 0.03 | 0.05 | 184.52 | 0.60 | .551 | [-0.07, 0.13] |
| ICPS frontolateral * Condition * Age | -0.03 | 0.06 | 340.23 | -0.55 | .583 | [-0.16, 0.09] |

*Note.* Significance codes: * p < .05, ** p < .01, *** p < .001. Degrees of freedom for the fixed effects were estimated using Satterthwaite's approximation.

**Table S9**

*Midfrontal-Midlateral ICPS Predicting SSP-DDM Boundary Separation (a) Model Statistics*

| Parameter | *β* | *SE* | *df* | *t* | *p* | 95% CI |
|---|---|---|---|---|---|---|
| (Intercept) | -0.02 | 0.08 | 203.46 | -0.22 | .826 | [-0.17, 0.14] |
| ICPS midlateral | 0.02 | 0.06 | 295.60 | 0.37 | .709 | [-0.09, 0.13] |
| Condition | -0.03 | 0.05 | 186.64 | -0.66 | .507 | [-0.14, 0.07] |

| | | | | | | |
|---|---|---|---|---|---|---|
| Age | 0.11 | 0.05 | 189.87 | 1.92 | .056 | [-0.00, 0.21] |
| Sex | 0.13 | 0.05 | 187.20 | 2.33 | .021* | [0.02, 0.23] |
| Peer mode | 0.05 | 0.08 | 201.10 | 0.57 | .567 | [-0.11, 0.20] |
| ICPS midlateral * Condition | -0.05 | 0.06 | 278.58 | -0.86 | .388 | [-0.16, 0.06] |
| ICPS midlateral * Age | -0.10 | 0.06 | 313.37 | -1.77 | .078 | [-0.21, 0.01] |
| Condition * Age | 0.03 | 0.05 | 188.74 | 0.65 | .519 | [-0.07, 0.14] |
| ICPS midlateral * Condition * Age | 0.05 | 0.06 | 299.39 | 0.90 | .370 | [-0.06, 0.16] |

*Note.* Significance codes: * p < .05, ** p < .01, *** p < .001. Degrees of freedom for the fixed effects were estimated using Satterthwaite's approximation.

**Table S10**

*Midfrontal-Posterolateral ICPS Predicting SSP-DDM Boundary Separation (a) Model Statistics*

| Parameter | *β* | *SE* | *df* | *t* | *p* | 95% CI |
|---|---|---|---|---|---|---|
| (Intercept) | -0.04 | 0.08 | 191.09 | -0.50 | .615 | [-0.19, 0.11] |
| ICPS posterolateral | 0.07 | 0.06 | 306.83 | 1.28 | .203 | [-0.04, 0.18] |
| Condition | -0.01 | 0.05 | 179.45 | -0.10 | .917 | [-0.11, 0.10] |
| Age | 0.10 | 0.05 | 177.13 | 1.95 | .053 | [0.00, 0.21] |
| Sex | 0.14 | 0.05 | 180.07 | 2.74 | .007** | [0.04, 0.25] |
| Peer mode | 0.07 | 0.08 | 192.13 | 0.86 | .388 | [-0.08, 0.21] |
| ICPS posterolateral * Condition | -0.03 | 0.06 | 297.68 | -0.57 | .569 | [-0.14, 0.08] |
| ICPS posterolateral * Age | 0.05 | 0.06 | 327.74 | 0.88 | .379 | [-0.06, 0.16] |
| Condition * Age | 0.06 | 0.05 | 177.23 | 1.11 | .270 | [-0.04, 0.16] |
| ICPS posterolateral * Condition * Age | -0.04 | 0.06 | 319.90 | -0.63 | .526 | [-0.15, 0.07] |

*Note.* Significance codes: * p < .05, ** p < .01, *** p < .001. Degrees of freedom for the fixed effects were estimated using Satterthwaite's approximation.

### Analyses with Raw Behavioral Measures of Post-Error Adjustments

For completeness, we also performed exploratory analyses examining relations between error-related theta ICPS and raw behavioral measures of post-error adjustments (PEA, PES, and PERI), as these measures were used in prior studies on the functional role of error-related theta ICPS (Buzzell et al., 2019; Cavanagh et al., 2009; Cavanagh & Frank, 2014). Using the same trial subset as for the DDM, we calculated three behavioral measures: (1) Post-error accuracy (PEA): accuracy difference between post-error and post-correct trials (greater values = better post-error performance); (2) Post-error slowing (PES): RT difference for correct post-error and post-correct trials (greater values = more RT slowing after errors); and (3) Post-error reduction in interference (PERI): difference in congruency effects (incongruent-congruent RT) between post-error and post-correct trials (smaller values = reduced post-error flanker interference). The set of LMMs we fit to each of the three measures was identical to the models with SSP-DDM measures, thus comprising a single model with three ICPS predictors and three follow-up models with each ICPS measure entered separately (Tables S11-S22). Below, we briefly report main effects and interactions involving ICPS in the full (three ICPS predictor) models.

Within the full, three-ICPS predictor models, we identified a significant main effect of midfrontal-posterolateral ICPS predicting higher PEA (β = 0.11, 95% CI [0.00, 0.21], $p$ = .046; Table S11). Additionally, we found that midfrontal-frontolateral theta ICPS predicted lower PES (β = -0.12, 95% CI [-0.23, -0.01], $p$ = .034; Table S19), while midfrontal-midlateral theta ICPS predicted higher PES (β = 0.14, 95% CI [0.02, 0.27], $p$ = .024; Table S19).

**Table S11**

*Full Error-Related Theta ICPS Predicting PEA Model Statistics*

| Parameter | *β* | *SE* | *df* | *t* | *p* | 95% CI |
|---|---|---|---|---|---|---|
| (Intercept) | -0.08 | 0.08 | 199.54 | -1.04 | .299 | [-0.23, 0.07] |
| ICPS frontolateral | -0.03 | 0.06 | 351.82 | -0.46 | .648 | [-0.14, 0.09] |
| Condition | -0.03 | 0.05 | 185.66 | -0.64 | .525 | [-0.13, 0.06] |
| Age | 0.16 | 0.05 | 194.71 | 2.87 | .005** | [0.05, 0.26] |
| ICPS midlateral | 0.02 | 0.06 | 333.85 | 0.33 | .744 | [-0.10, 0.14] |
| ICPS posterolateral | 0.11 | 0.05 | 346.92 | 2.00 | .046* | [0.00, 0.21] |
| Sex | 0.07 | 0.05 | 186.33 | 1.36 | .174 | [-0.03, 0.17] |
| Peer mode | 0.08 | 0.08 | 199.03 | 1.05 | .294 | [-0.07, 0.23] |
| ICPS frontolateral * Condition | 0.02 | 0.06 | 336.90 | 0.41 | .681 | [-0.09, 0.14] |
| ICPS frontolateral * Age | -0.03 | 0.06 | 353.44 | -0.44 | .657 | [-0.14, 0.09] |
| Condition * Age | 0.03 | 0.05 | 187.67 | 0.63 | .532 | [-0.06, 0.13] |
| Condition * ICPS midlateral | 0.06 | 0.06 | 297.77 | 0.98 | .328 | [-0.06, 0.18] |
| Age * ICPS midlateral | 0.01 | 0.06 | 341.42 | 0.11 | .914 | [-0.11, 0.12] |
| Condition * ICPS posterolateral | 0.03 | 0.05 | 322.76 | 0.56 | .576 | [-0.07, 0.13] |
| Age * ICPS posterolateral | 0.09 | 0.05 | 353.99 | 1.63 | .105 | [-0.02, 0.19] |
| ICPS frontolateral * Condition * Age | -0.03 | 0.06 | 344.96 | -0.55 | .581 | [-0.14, 0.08] |
| Condition * Age * ICPS midlateral | -0.03 | 0.06 | 315.76 | -0.55 | .582 | [-0.14, 0.08] |
| Condition * Age * ICPS posterolateral | -0.03 | 0.05 | 345.48 | -0.61 | .542 | [-0.13, 0.07] |

*Note.* Significance codes: * p < .05, ** p < .01, *** p < .001. Degrees of freedom for the fixed effects were estimated using Satterthwaite's approximation.

**Table S12**

*Midfrontal-Frontolateral Theta ICPS Predicting PEA Model Statistics*

| Parameter | *β* | *SE* | *df* | *t* | *p* | 95% CI |
|---|---|---|---|---|---|---|
| (Intercept) | -0.08 | 0.08 | 216.45 | -0.96 | .336 | [-0.23, 0.08] |
| ICPS frontolateral | 0.03 | 0.05 | 385.78 | 0.50 | .619 | [-0.07, 0.13] |
| Condition | -0.03 | 0.05 | 194.56 | -0.64 | .526 | [-0.12, 0.06] |
| Age | 0.18 | 0.05 | 205.12 | 3.27 | .001** | [0.07, 0.28] |
| Sex | 0.04 | 0.05 | 197.36 | 0.72 | .474 | [-0.06, 0.14] |
| Peer mode | 0.08 | 0.08 | 215.78 | 1.08 | .283 | [-0.07, 0.23] |
| ICPS frontolateral * Condition | 0.03 | 0.05 | 347.67 | 0.62 | .532 | [-0.07, 0.13] |
| ICPS frontolateral * Age | -0.05 | 0.05 | 388.86 | -1.01 | .314 | [-0.16, 0.05] |
| Condition * Age | 0.03 | 0.05 | 197.78 | 0.64 | .521 | [-0.06, 0.12] |
| ICPS frontolateral * Condition * Age | -0.06 | 0.05 | 364.86 | -1.08 | .281 | [-0.16, 0.05] |

*Note.* Significance codes: * p < .05, ** p < .01, *** p < .001. Degrees of freedom for the fixed effects were estimated using Satterthwaite's approximation.

**Table S13**

*Midfrontal-Midlateral Theta ICPS Predicting PEA Model Statistics*

| Parameter | *β* | *SE* | *df* | *t* | *p* | 95% CI |
|---|---|---|---|---|---|---|
| (Intercept) | -0.06 | 0.08 | 214.74 | -0.74 | .457 | [-0.20, 0.09] |
| ICPS midlateral | 0.07 | 0.05 | 371.30 | 1.38 | .170 | [-0.03, 0.17] |
| Condition | -0.04 | 0.05 | 208.43 | -0.78 | .437 | [-0.13, 0.05] |
| Age | 0.16 | 0.05 | 219.15 | 3.00 | .003** | [0.06, 0.26] |
| Sex | 0.05 | 0.05 | 210.71 | 0.89 | .377 | [-0.06, 0.15] |
| Peer mode | 0.07 | 0.08 | 210.61 | 0.94 | .348 | [-0.08, 0.22] |
| ICPS midlateral * Condition | 0.05 | 0.05 | 312.50 | 1.06 | .289 | [-0.04, 0.15] |
| ICPS midlateral * Age | -0.03 | 0.05 | 383.87 | -0.50 | .614 | [-0.12, 0.07] |
| Condition * Age | 0.01 | 0.05 | 212.88 | 0.12 | .908 | [-0.08, 0.09] |
| ICPS midlateral * Condition * Age | -0.02 | 0.05 | 340.86 | -0.45 | .650 | [-0.12, 0.07] |

*Note.* Significance codes: * p < .05, ** p < .01, *** p < .001. Degrees of freedom for the fixed effects were estimated using Satterthwaite's approximation.

**Table S14**

*Midfrontal-Posterolateral Theta ICPS Predicting PEA Model Statistics*

| Parameter | *β* | *SE* | *df* | *t* | *p* | 95% CI |
|---|---|---|---|---|---|---|
| (Intercept) | -0.07 | 0.07 | 200.87 | -0.96 | .337 | [-0.21, 0.07] |
| ICPS posterolateral | 0.10 | 0.05 | 369.00 | 2.01 | .045* | [0.00, 0.20] |
| Condition | -0.05 | 0.05 | 199.93 | -1.09 | .279 | [-0.14, 0.04] |

| | | | | | | |
|---|---|---|---|---|---|---|
| Age | 0.16 | 0.05 | 208.02 | 3.15 | .002** | [0.06, 0.26] |
| Sex | 0.05 | 0.05 | 203.26 | 1.01 | .312 | [-0.05, 0.15] |
| Peer mode | 0.07 | 0.07 | 200.42 | 1.01 | .313 | [-0.07, 0.21] |
| ICPS posterolateral * Condition | 0.04 | 0.05 | 329.40 | 0.92 | .356 | [-0.05, 0.14] |
| ICPS posterolateral * Age | 0.10 | 0.05 | 387.55 | 2.00 | .046* | [0.00, 0.19] |
| Condition * Age | 0.04 | 0.05 | 203.68 | 0.90 | .367 | [-0.05, 0.13] |
| ICPS posterolateral * Condition * Age | -0.03 | 0.05 | 367.24 | -0.70 | .485 | [-0.13, 0.06] |

*Note.* Significance codes: * p < .05, ** p < .01, *** p < .001. Degrees of freedom for the fixed effects were estimated using Satterthwaite's approximation.

**Table S15**

*Full Error-Related Theta ICPS Predicting PERI Model Statistics*

| Parameter | *β* | *SE* | *df* | *t* | *p* | 95% CI |
|---|---|---|---|---|---|---|
| (Intercept) | 0.03 | 0.08 | 213.35 | 0.43 | .671 | [-0.12, 0.18] |
| ICPS frontolateral | 0.11 | 0.06 | 350.67 | 1.73 | .085 | [-0.01, 0.22] |
| Condition | 0.10 | 0.05 | 198.35 | 1.96 | .051 | [0.00, 0.20] |
| Age | -0.01 | 0.05 | 206.28 | -0.19 | .848 | [-0.12, 0.09] |
| ICPS midlateral | -0.12 | 0.06 | 335.70 | -1.85 | .065 | [-0.25, 0.00] |
| ICPS posterolateral | 0.02 | 0.06 | 344.75 | 0.28 | .782 | [-0.09, 0.12] |
| Sex | -0.03 | 0.05 | 197.40 | -0.49 | .621 | [-0.13, 0.08] |
| Peer mode | 0.00 | 0.08 | 212.87 | 0.03 | .979 | [-0.15, 0.15] |
| ICPS frontolateral * Condition | -0.00 | 0.06 | 340.52 | -0.00 | .999 | [-0.12, 0.12] |
| ICPS frontolateral * Age | -0.01 | 0.06 | 352.46 | -0.19 | .852 | [-0.13, 0.10] |
| Condition * Age | 0.03 | 0.05 | 201.16 | 0.61 | .546 | [-0.07, 0.13] |
| Condition * ICPS midlateral | -0.02 | 0.06 | 309.89 | -0.26 | .793 | [-0.14, 0.10] |
| Age * ICPS midlateral | -0.01 | 0.06 | 343.14 | -0.11 | .914 | [-0.12, 0.11] |
| Condition * ICPS posterolateral | 0.02 | 0.05 | 327.11 | 0.34 | .734 | [-0.09, 0.12] |
| Age * ICPS posterolateral | 0.03 | 0.05 | 352.97 | 0.53 | .599 | [-0.08, 0.13] |
| ICPS frontolateral * Condition * Age | 0.03 | 0.06 | 346.81 | 0.43 | .665 | [-0.09, 0.14] |
| Condition * Age * ICPS midlateral | -0.07 | 0.06 | 326.75 | -1.25 | .213 | [-0.19, 0.04] |
| Condition * Age * ICPS posterolateral | 0.07 | 0.05 | 346.85 | 1.32 | .188 | [-0.03, 0.17] |

*Note.* Significance codes: * p < .05, ** p < .01, *** p < .001. Degrees of freedom for the fixed effects were estimated using Satterthwaite's approximation.

**Table S16**

*Midfrontal-Frontolateral Theta ICPS Predicting PERI Model Statistics*

| Parameter | *β* | *SE* | *df* | *t* | *p* | 95% CI |
|---|---|---|---|---|---|---|
| (Intercept) | 0.02 | 0.08 | 226.82 | 0.21 | .836 | [-0.14, 0.17] |
| ICPS frontolateral | 0.03 | 0.05 | 379.31 | 0.66 | .512 | [-0.07, 0.14] |
| Condition | 0.11 | 0.05 | 205.52 | 2.25 | .025* | [0.02, 0.21] |

| | | | | | | |
|---|---|---|---|---|---|---|
| Age | -0.02 | 0.05 | 213.66 | -0.46 | .647 | [-0.13, 0.08] |
| Sex | -0.01 | 0.05 | 204.08 | -0.13 | .896 | [-0.11, 0.09] |
| Peer mode | -0.02 | 0.08 | 225.44 | -0.32 | .749 | [-0.18, 0.13] |
| ICPS frontolateral * Condition | -0.00 | 0.05 | 359.52 | -0.07 | .945 | [-0.11, 0.10] |
| ICPS frontolateral * Age | 0.01 | 0.06 | 385.26 | 0.10 | .916 | [-0.10, 0.11] |
| Condition * Age | 0.06 | 0.05 | 210.22 | 1.12 | .266 | [-0.04, 0.15] |
| ICPS frontolateral * Condition * Age | 0.00 | 0.05 | 372.42 | 0.05 | .957 | [-0.10, 0.11] |

*Note.* Significance codes: * p < .05, ** p < .01, *** p < .001. Degrees of freedom for the fixed effects were estimated using Satterthwaite's approximation.

**Table S17**

*Midfrontal-Midlateral Theta ICPS Predicting PERI Model Statistics*

| Parameter | *β* | *SE* | *df* | *t* | *p* | 95% CI |
|---|---|---|---|---|---|---|
| (Intercept) | 0.02 | 0.07 | 211.09 | 0.30 | .767 | [-0.12, 0.17] |
| ICPS midlateral | -0.04 | 0.05 | 349.15 | -0.77 | .442 | [-0.14, 0.06] |
| Condition | 0.11 | 0.05 | 208.79 | 2.23 | .027* | [0.01, 0.21] |
| Age | -0.01 | 0.05 | 216.77 | -0.15 | .883 | [-0.11, 0.09] |
| Sex | 0.01 | 0.05 | 206.84 | 0.10 | .921 | [-0.09, 0.10] |
| Peer mode | -0.02 | 0.07 | 208.46 | -0.24 | .810 | [-0.16, 0.13] |
| ICPS midlateral * Condition | -0.05 | 0.05 | 329.48 | -0.99 | .323 | [-0.15, 0.05] |
| ICPS midlateral * Age | -0.01 | 0.05 | 370.57 | -0.28 | .783 | [-0.12, 0.09] |
| Condition * Age | 0.05 | 0.05 | 215.89 | 1.00 | .318 | [-0.05, 0.15] |
| ICPS midlateral * Condition * Age | -0.02 | 0.05 | 358.72 | -0.44 | .659 | [-0.12, 0.08] |

*Note.* Significance codes: * p < .05, ** p < .01, *** p < .001. Degrees of freedom for the fixed effects were estimated using Satterthwaite's approximation.

**Table S18**

*Midfrontal-Posterolateral Theta ICPS Predicting PERI Model Statistics*

| Parameter | *β* | *SE* | *df* | *t* | *p* | 95% CI |
|---|---|---|---|---|---|---|
| (Intercept) | 0.05 | 0.07 | 202.55 | 0.65 | .517 | [-0.09, 0.19] |
| ICPS posterolateral | -0.01 | 0.05 | 362.14 | -0.18 | .860 | [-0.11, 0.09] |
| Condition | 0.08 | 0.05 | 200.17 | 1.63 | .105 | [-0.02, 0.17] |
| Age | -0.02 | 0.05 | 206.58 | -0.45 | .655 | [-0.12, 0.08] |
| Sex | -0.02 | 0.05 | 201.95 | -0.32 | .751 | [-0.11, 0.08] |
| Peer mode | -0.03 | 0.07 | 202.25 | -0.42 | .676 | [-0.17, 0.11] |
| ICPS posterolateral * Condition | -0.01 | 0.05 | 333.08 | -0.26 | .791 | [-0.11, 0.08] |
| ICPS posterolateral * Age | 0.02 | 0.05 | 384.54 | 0.37 | .712 | [-0.08, 0.12] |
| Condition * Age | 0.03 | 0.05 | 204.24 | 0.67 | .505 | [-0.06, 0.12] |
| ICPS posterolateral * Condition * Age | 0.04 | 0.05 | 369.32 | 0.88 | .379 | [-0.05, 0.14] |

*Note.* Significance codes: * p < .05, ** p < .01, *** p < .001. Degrees of freedom for the fixed effects were estimated using Satterthwaite's approximation.

**Table S19**

*Full Error-Related Theta ICPS Predicting PES Model Statistics*

| Parameter | *β* | *SE* | *df* | *t* | *p* | 95% CI |
|---|---|---|---|---|---|---|
| (Intercept) | -0.01 | 0.08 | 212.51 | -0.15 | .879 | [-0.17, 0.14] |
| ICPS frontolateral | -0.12 | 0.06 | 353.27 | -2.13 | .034* | [-0.23, -0.01] |
| Condition | -0.04 | 0.05 | 185.47 | -0.79 | .432 | [-0.12, 0.05] |
| Age | 0.04 | 0.06 | 205.78 | 0.71 | .478 | [-0.07, 0.15] |
| ICPS midlateral | 0.14 | 0.06 | 349.52 | 2.27 | .024* | [0.02, 0.27] |
| ICPS posterolateral | -0.04 | 0.05 | 353.94 | -0.67 | .502 | [-0.14, 0.07] |
| Sex | 0.24 | 0.05 | 197.46 | 4.31 | < .001*** | [0.13, 0.34] |
| Peer mode | 0.06 | 0.08 | 210.36 | 0.72 | .475 | [-0.09, 0.21] |
| ICPS frontolateral * Condition | 0.03 | 0.06 | 320.96 | 0.61 | .545 | [-0.07, 0.14] |
| ICPS frontolateral * Age | -0.06 | 0.06 | 352.95 | -1.07 | .286 | [-0.17, 0.05] |
| Condition * Age | 0.12 | 0.04 | 186.96 | 2.60 | .010** | [0.03, 0.20] |
| Condition * ICPS midlateral | -0.10 | 0.06 | 275.43 | -1.68 | .095 | [-0.21, 0.01] |
| Age * ICPS midlateral | 0.02 | 0.06 | 351.44 | 0.26 | .797 | [-0.10, 0.13] |
| Condition * ICPS posterolateral | -0.03 | 0.05 | 301.36 | -0.54 | .586 | [-0.12, 0.07] |
| Age * ICPS posterolateral | -0.06 | 0.05 | 348.89 | -1.16 | .247 | [-0.16, 0.04] |
| ICPS frontolateral * Condition * Age | -0.05 | 0.06 | 333.38 | -0.83 | .409 | [-0.15, 0.06] |
| Condition * Age * ICPS midlateral | 0.04 | 0.05 | 295.64 | 0.73 | .467 | [-0.06, 0.14] |
| Condition * Age * ICPS posterolateral | 0.07 | 0.05 | 328.94 | 1.33 | .183 | [-0.03, 0.16] |

*Note.* Significance codes: * p < .05, ** p < .01, *** p < .001. Degrees of freedom for the fixed effects were estimated using Satterthwaite's approximation.

**Table S20**

*Midfrontal-Frontolateral Theta ICPS Predicting PES Model Statistics*

| Parameter | *β* | *SE* | *df* | *t* | *p* | 95% CI |
|---|---|---|---|---|---|---|
| (Intercept) | -0.02 | 0.08 | 229.19 | -0.25 | .802 | [-0.17, 0.13] |
| ICPS frontolateral | -0.07 | 0.05 | 388.46 | -1.48 | .139 | [-0.17, 0.02] |
| Condition | -0.03 | 0.04 | 197.70 | -0.71 | .480 | [-0.11, 0.05] |
| Age | 0.07 | 0.05 | 215.61 | 1.20 | .233 | [-0.04, 0.17] |
| Sex | 0.24 | 0.05 | 208.08 | 4.37 | < .001*** | [0.13, 0.34] |
| Peer mode | 0.05 | 0.08 | 228.38 | 0.65 | .519 | [-0.10, 0.21] |
| ICPS frontolateral * Condition | 0.01 | 0.05 | 325.43 | 0.12 | .903 | [-0.09, 0.10] |
| ICPS frontolateral * Age | -0.05 | 0.05 | 385.51 | -1.03 | .302 | [-0.15, 0.05] |
| Condition * Age | 0.11 | 0.04 | 200.79 | 2.54 | .012* | [0.03, 0.19] |
| ICPS frontolateral * Condition * Age | -0.02 | 0.05 | 345.33 | -0.34 | .733 | [-0.11, 0.08] |

*Note.* Significance codes: * p < .05, ** p < .01, *** p < .001. Degrees of freedom for the fixed effects were estimated using Satterthwaite's approximation.

**Table S21**

*Midfrontal-Midlateral Theta ICPS Predicting PES Model Statistics*

| Parameter | *β* | *SE* | *df* | *t* | *p* | 95% CI |
|---|---|---|---|---|---|---|
| (Intercept) | -0.06 | 0.08 | 220.55 | -0.85 | .398 | [-0.21, 0.08] |
| ICPS midlateral | 0.04 | 0.05 | 376.21 | 0.75 | .452 | [-0.06, 0.14] |
| Condition | -0.05 | 0.04 | 210.23 | -1.11 | .268 | [-0.13, 0.04] |
| Age | 0.01 | 0.05 | 222.31 | 0.22 | .828 | [-0.09, 0.11] |
| Sex | 0.23 | 0.05 | 214.18 | 4.48 | < .001*** | [0.13, 0.33] |
| Peer mode | 0.08 | 0.07 | 216.12 | 1.05 | .293 | [-0.07, 0.22] |
| ICPS midlateral * Condition | -0.07 | 0.05 | 308.02 | -1.49 | .137 | [-0.16, 0.02] |
| ICPS midlateral * Age | 0.00 | 0.05 | 386.39 | 0.10 | .920 | [-0.09, 0.10] |
| Condition * Age | 0.12 | 0.04 | 213.87 | 2.77 | .006** | [0.04, 0.21] |
| ICPS midlateral * Condition * Age | 0.05 | 0.05 | 335.32 | 1.12 | .262 | [-0.04, 0.14] |

*Note.* Significance codes: * p < .05, ** p < .01, *** p < .001. Degrees of freedom for the fixed effects were estimated using Satterthwaite's approximation.

**Table S22**

*Midfrontal-Posterolateral Theta ICPS Predicting PES Model Statistics*

| Parameter | *β* | *SE* | *df* | *t* | *p* | 95% CI |
|---|---|---|---|---|---|---|
| (Intercept) | -0.04 | 0.07 | 207.15 | -0.54 | .590 | [-0.19, 0.10] |
| ICPS posterolateral | -0.03 | 0.05 | 382.14 | -0.59 | .556 | [-0.12, 0.07] |
| Condition | -0.03 | 0.04 | 197.32 | -0.83 | .410 | [-0.12, 0.05] |
| Age | 0.04 | 0.05 | 211.02 | 0.76 | .449 | [-0.06, 0.14] |
| Sex | 0.24 | 0.05 | 206.87 | 4.54 | < .001*** | [0.14, 0.34] |
| Peer mode | 0.07 | 0.07 | 206.54 | 0.89 | .376 | [-0.08, 0.21] |
| ICPS posterolateral * Condition | -0.03 | 0.05 | 308.34 | -0.70 | .487 | [-0.12, 0.06] |
| ICPS posterolateral * Age | -0.05 | 0.05 | 386.37 | -1.09 | .279 | [-0.14, 0.04] |
| Condition * Age | 0.09 | 0.04 | 200.22 | 2.21 | .028* | [0.01, 0.18] |
| ICPS posterolateral * Condition * Age | 0.09 | 0.05 | 347.99 | 2.03 | .043* | [0.00, 0.18] |

*Note.* Significance codes: * p < .05, ** p < .01, *** p < .001. Degrees of freedom for the fixed effects were estimated using Satterthwaite's approximation.
